# Targeting DNA-LNPs to Endothelial Cells Improves Expression Magnitude, Duration, and Specificity

**DOI:** 10.1101/2025.07.09.663747

**Authors:** Nicolas Marzolini, Taylor V. Brysgel, Ryan J. Rahman, Eno-Obong Essien, Saw Y. Nwe, Jichuan Wu, Aparajeeta Majumder, Manthan N. Patel, Sachchidanand Tiwari, Carolann L. Espy, Fengyi Dong, Anit Shah, Vladimir V. Shuvaev, Elizabeth D. Hood, Liam S. Chase, Drew Weissman, Jeremy B. Katzen, David B. Frank, Mariko L. Bennett, Oscar A. Marcos-Contreras, Jacob W. Myerson, Vladimir R. Muzykantov, Sahily Reyes-Esteves, Jacob S. Brenner

## Abstract

DNA-lipid nanoparticles (DNA-LNPs) loaded with inhibitors of the cGAS-STING pathway enable safe and effective delivery of DNA *in vivo*. Herein, we report the first instances of extrahepatic DNA-LNP targeting. DNA-LNPs conjugated to antibodies against PECAM-1 or VCAM-1 target the endothelium of the lungs and brain/spleen, respectively. These LNPs drive robust transgene expression in their target organs, with greater magnitude and duration than untargeted LNPs. Lung specificity of PECAM-targeted transgene expression increases over two weeks, resulting in markedly higher lung-to-liver expression ratios than our previous PECAM-targeted mRNA-LNPs. Off-target liver DNA expression declines to undetectable levels but persists in the lungs, while mRNA expression uniformly decreases due to its short half-life. We further improve this expression specificity by replacing full-length antibodies with Fab fragments. Single-cell analysis reveals a key mechanism underlying the improvements in organ-specificity: target organ expression is dominated by long-lived endothelial cells, while off-target liver delivery and expression are in non-endothelial cells with shorter half-lives. Collectively, these studies demonstrate that targeted DNA-LNPs achieve high levels of organ- and cell-type-specific transgene expression and thus provide a therapeutic platform for dozens of endothelial-centric diseases.

## 1. Introduction

Lipid nanoparticles (LNPs) are a versatile delivery system with immense therapeutic potential due to their ability to encapsulate nucleic acid therapies^1^. In the past, LNPs have traditionally been loaded with messenger RNA (mRNA), as in the COVID-19 vaccines, to express encoded proteins as therapies or antigens^2^. LNPs loaded with small interfering RNAs (siRNAs) to silence gene expression have also seen therapeutic success, as Patisiran was the first FDA-approved lipid nanoparticle^3^. Despite these accomplishments, RNA-LNPs have not yet been clinically translated for chronic indications due to the relatively short half-life of RNA^1^. Current literature is full of examples using RNA-LNPs to treat chronic diseases via the delivery of CRISPR/Cas9 machinery, base editors, siRNA, and other gene editing machinery^4,5^. However, these approaches raise concerns regarding off-target editing and are limited to chronic diseases with well-defined genetic mutations^6^. Therefore, DNA-LNPs are an attractive drug delivery system based on their potential for long-term expression, less frequent dosing, and enhanced stability^7^.

Historically, DNA-LNPs have been too inflammatory to use *in vivo*, as the delivery of DNA to the cytosol activates the cyclic GMP-AMP synthase/stimulator of interferon genes (cGAS–STING) signaling pathway, leading to the upregulation of pro-inflammatory cytokines and type 1 interferons that induce 100% mortality at biologically relevant doses^7–9^. Our lab has recently published methods to reduce this inflammation by co-loading DNA-LNPs with 9(10)-nitrooleic acid (NOA), a lipid inhibitor of STING^9,10^. These NOA-DNA-LNPs significantly alleviate inflammation *in vivo* and reduce mortality to 0% at doses up to 25µg of plasmid DNA (pDNA) loaded into bare (unconjugated) LNPs^9^. Further, NOA-DNA-LNPs express > 6 months per dose, have much larger cargo capacity than the dominant viral vector (adeno-associated virus, or AAV), and can be redosed^11,12^. With these characteristics, DNA-LNPs are poised to fill niches in gene therapy which cannot be served by AAVs and mRNA-LNPs, such as delivery of large transgenes, delivery to patients with anti-AAV antibodies, and treatment of chronic diseases without a clear, single genetic origin (*e.g.*, atherosclerosis, osteoarthritis, etc.)^13–15^.

When administered intravenously (IV), most unmodified LNPs, whether carrying DNA or RNA, accumulate predominantly in the liver^16^. This makes it difficult to use LNPs to treat most diseases outside the liver. Therefore, we started the campaign to target DNA-LNPs to specific organs and cell types. As our first target, we chose endothelial cells, for two reasons. First, endothelial cells play a central role in many illnesses, including pulmonary hypertension (PH) and stroke^17,18^. Second, if we can target DNA-LNPs to capillary endothelial cells in a specific organ, we could turn those cells into biofactories which secrete therapeutic proteins into the parenchyma of that organ, a strategy that could treat many diseases that are not endothelial-centric^19^.

Historically, two approaches to achieve targeted delivery of LNPs to the endothelium have been implemented: (1) attaching affinity ligands, such as antibodies against cell-specific targets, to their surfaces or (2) modifying the physical or chemical features of the LNP, such as introducing permanently cationic lipids like 1,2-dioleoyl-3-trimethylammonium propane (DOTAP) into the formulation^20–23^. However, physicochemical tropism has been shown to induce severe toxicities in vivo (*e.g.*, thrombosis)^24^. We thus utilize affinity targeting of endothelial cell surface proteins. Platelet endothelial cell adhesion molecule 1 (PECAM-1) is an attractive candidate for redirecting the delivery of LNPs to the lungs, since it is highly expressed on the surface of pulmonary endothelial cells^25^. Moreover, the lungs receive the entire cardiac output and have a large vascular surface area^26^. Conjugating anti-PECAM-1 (αPECAM) antibodies to the surface of mRNA-LNPs has been shown to redirect their delivery to the lungs, achieving ∼100% injected dose per gram tissue (%ID/g)^27^. Similarly, conjugating antibodies against vascular cell adhesion molecule 1 (VCAM-1), another adhesion molecule whose expression is upregulated in inflammatory conditions, increases the delivery and expression of targeted nanocarriers to the endothelium of the spleen and brain^28–32^.

Here, we adapt this strategy for NOA-DNA-LNPs (referred to as DNA-LNPs, hereafter) to target them to the endothelium of different organs. By conjugating monoclonal antibodies (mAbs) against PECAM-1 or VCAM-1 to nanoparticles, we redirect their delivery and expression to the lungs and spleen/brain, respectively. *Ex vivo* imaging of organ luminescence confirms this DNA expression as being concentrated in the target organs and demonstrates that the lung versus liver specificity of our targeted DNA-LNP expression increases over time and exceeds that of targeted mRNA-LNPs. Replacing full-length PECAM mAbs with fragment-antigen binding regions (Fabs) further increases the magnitude, longevity, and organ-type specificity of transgene expression. Using flow cytometry and immunofluorescence, we confirm pulmonary endothelial cells as the primary cell type targeted by our αPECAM-Fab nanoparticles and responsible for transgene expression. Overall, these results highlight the immense potential of targeted DNA-LNPs for treating a variety of chronic endothelial pathologies beyond the scope of traditional mRNA-LNPs and other gene therapies.

## 2. Results

### 2.1. Conjugating **α**PECAM antibodies to DNA-LNPs redirects their delivery to the lungs

We first fabricated DNA-LNPs based on Pfizer-BioNTech’s FDA-approved COVID-19 mRNA vaccine LNP formulation, which consists of the ionizable lipid ALC-0315, the phospholipid 1,2-distearoyl-sn-glycero-3-phosphocholine (DSPC), cholesterol, and ALC-0159, which was replaced fully with 1,2-distearoyl-sn-glycero-3-phosphoethanolamine -N- [azido(polyethylene glycol)-2000] (DSPE-PEG(2000)-Azide)^33^. Our LNPs were loaded with reporter plasmid DNA encoding for either luciferase, mCherry, or Cre recombinase (lipid/pDNA ratio of 40/1, wt/wt) (**Figure 1a**). The replacement of ALC-0159 with DSPE-PEG(2000)-Azide allowed us to conjugate antibodies to the surface of LNPs using established click chemistry methods^30,34^. Additionally, we co-loaded NOA into our lipid formulation, which inhibits the cytosolic DNA sensor, STING, thereby reducing inflammation and mitigating mortality as described in our prior work^9^. αPECAM monoclonal antibodies conjugated to DNA-LNPs with greater than 80% conjugation efficiency (**Figure S1)**, resulting in average size increases of 15-20nm post-conjugation and polydispersity indices (PDIs) consistently < 0.2 (**Figure 1b**).

**Figure 1.**
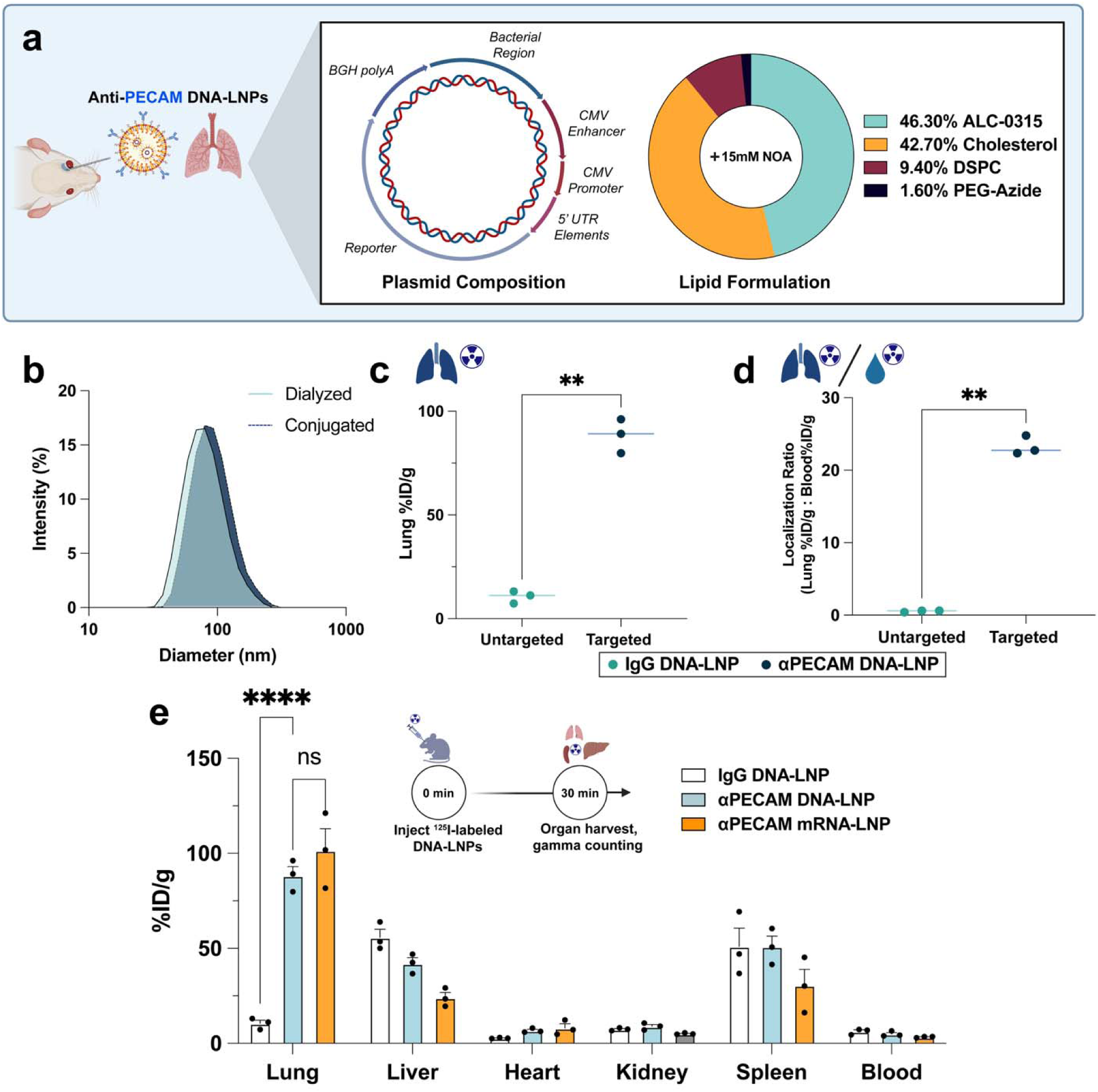
Anti-PECAM DNA-LNPs specifically target the lungs. (a) Graphical schematic depicting (left) treatment paradigm with retro-orbital injections of αPECAM DNA-LNPs with the illustrated lipid formulation and Nanoplasmid DNA. (b) DNA-LNP hydrodynamic diameter distribution before and after conjugation with αPECAM monoclonal antibodies as determined by dynamic light scattering (DLS). (c) Biodistribution study via radiotracing of ^125^I-labeled DNA-LNPs reveals ∼9-fold increase in lung uptake of αPECAM DNA-LNPs compared to untargeted IgG DNA-LNPs. %ID/g represents the percent of total injected dose detected in each organ normalized by respective organ mass. (d) Localization ratios accounting for blood retention of DNA-LNPs display ∼35x higher lung uptake of αPECAM DNA-LNPs compared with untargeted IgG controls when normalizing by blood signal. (e) Biodistribution of major abdominal and thoracic organs shows highest nanoparticle uptake in the lungs of mice administered αPECAM DNA- or mRNA-LNPs with no statistically significant difference in any other measured organ. For panels (c) and (d), unpaired *t*-tests with Welch’s correction were performed, and lines are drawn through median. In panel (e), a two-way ANOVA with Tukey’s multiple comparisons test was performed. All data include n = 3 and represent mean ± SEM; ** = *p* < 0.01, **** = *p* < 0.0001, ns = not significant.

For biodistribution studies, we intravenously injected naive, BALB/c mice with either (1) DNA- or mRNA-LNPs conjugated to αPECAM mAbs (∼50 per particle) and radiolabeled, untargeted immunoglobulin G (IgG, 5 mAbs per particle) to track particle distribution or (2) DNA-LNPs conjugated to untargeted IgG and radiolabeled IgG (∼50 unmodified IgG, 5 radiolabeled IgG per particle). Each mouse was dosed by total injected radioactivity in counts per minute (cpm) that equated to ∼2.5μg of nucleic acid. 30 minutes after injection, all mice were euthanized to determine the organ distribution of the particles. αPECAM DNA-LNPs showed increased lung accumulation compared to untargeted IgG DNA-LNP controls, with a percent injected dose per gram of tissue (%ID/g) of ∼90% (**Figure 1c**), underscoring that targeting PECAM-1 effectively redirects the delivery of LNPs to the lungs. Notably, αPECAM DNA-LNPs had no significant difference in delivery to any organ other than the lungs, although delivery to the liver downtrends (**Figure 1e**). Likewise, αPECAM DNA-LNPs target the lungs as well as αPECAM mRNA-LNPs, with no significant differences in the biodistribution lung-to-liver ratios between mRNA- and DNA-LNPs on a mouse-by-mouse basis (**Figure S3**). The localization ratio accounts for blood retention (tissue %ID/g normalized by whole blood %ID/g) and was ∼35x higher in αPECAM DNA-LNPs than IgG controls (**Figure 1d**). We also screened additional formulations of αPECAM DNA-LNPs using the ionizable lipid SM-102 and the phospholipid 1,2-Dioleoyl-sn-glycero-3-phosphoethanolamine (DOPE). While these alternative DNA-LNPs targeted the lungs similarly, our initial ALC-0315/DSPC formulation exhibited the highest conjugation efficiency to αPECAM mAbs (**Figure S1**) and matched the highest levels of lung targeting (**Figure S2**). Accordingly, our studies in the remainder of this paper utilize this LNP formulation.

### 2.2 áPECAM DNA-LNPs express in the lungs *in vivo*

We next investigated the transgene expression of our targeted αPECAM DNA-LNPs by measuring both *in vivo* and *ex vivo* luminescence. Initially, two doses were tested in naive mice with either 2.5 or 5μg of luciferase pDNA loaded into LNPs conjugated to αPECAM mAbs. The magnitude of *in vivo* luciferase expression in the thoracic cavity of the mice was measured using an in vivo imaging system (IVIS), which quantifies luminescence following intraperitoneal (IP) administration of the luciferase substrate, luciferin. One day following injection, luminescence in the αPECAM DNA-LNP-treated mice was visibly concentrated in the thorax, with highest levels of signal appearing to emit from the lungs (**Figures 2a, S4**). Total mean luminescence of mice treated with 5μg of pDNA reached a peak of ∼5 × 10^6^ photons s^-1^ at day 1, before steadily declining and plateauing around 7 × 10^5^ photons s^-1^ by day 7, which was stable at day 14, after which animals were euthanized for *ex vivo* analysis (**Figure 2c**). The day 14 luminescence in mice treated with αPECAM DNA-LNPs was significantly higher than that from untreated control mice (∼10^5^ photons s^-1^). Untargeted IgG and bare DNA-LNP controls (5μg pDNA) quickly plateaued by day 2, maintaining luminescence ∼2x greater than untreated controls yet substantially smaller than αPECAM DNA-LNPs thereafter (**Figure S5**). Furthermore, luminescence from mice treated with 5μg of pDNA loaded in αPECAM DNA-LNPs was higher than mice treated with 2.5μg of pDNA, whose signal was also concentrated in the lungs (**Figure S4**) and followed a similar trend of peaking within the first few days and plateauing by day 7 (**Figure 2c**).

**Figure 2.**
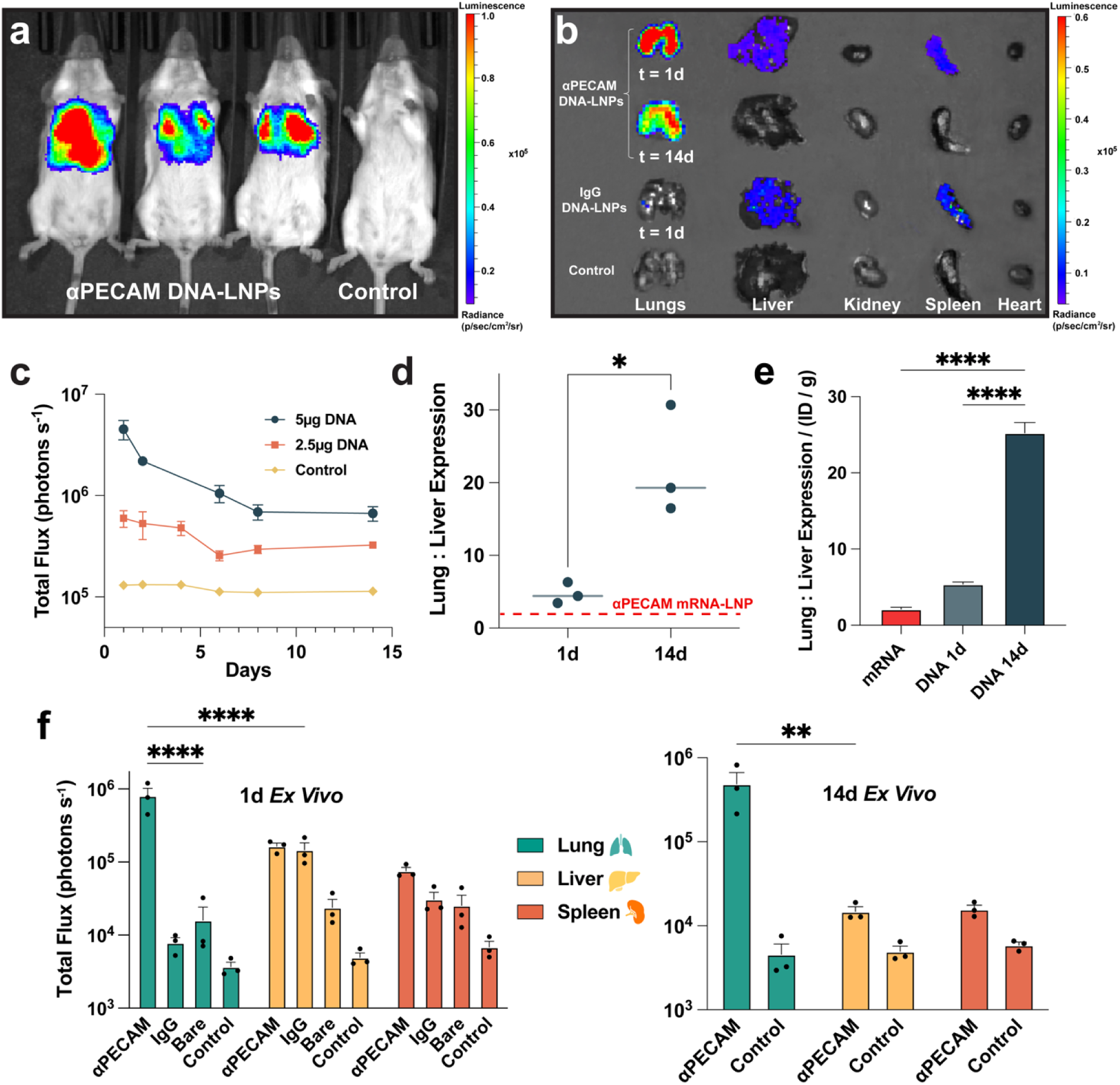
PECAM-targeted DNA-LNPs express in the lungs with increasing lung specificity over time. (a) Representative bioluminescence (IVIS) image of BALB/c mice treated with 5μg of luciferase pDNA loaded into αPECAM DNA-LNPs alongside a naive, luciferin-treated control shows a strong luminescent signal in the thoracic region, and notably the lobes of the lungs, at 1d. (b) Representative *ex vivo* IVIS image of major thoracic and abdominal organs from mice treated with αPECAM DNA-LNPs or IgG DNA-LNPs next to control organs from a naive, luciferin-treated mouse. Notably, luminescence is retained in the lungs at 14d and is greatly diminished in the liver. (c) Quantified total luminescence (represented as total flux in photons s^-1^) in the area of visible signal from *in vivo* IVIS images over time for mice treated with 5μg and 2.5μg of pDNA loaded into αPECAM DNA-LNPs compared to naive, luciferin-treated control. The expression of luciferase is dose-dependent and plateaus by day 7. (d) Lung-to-liver ratios of *ex vivo* luminescence underscore increasing lung specificity of expression over time from day 1 to day 14. Dashed line denotes the peak lung-to-liver ratio of αPECAM mRNA-LNPs previously reported by our group, where peak mRNA expression occurs at 4.5h post-injection. (e) Comparison of lung-to-liver expression ratios normalized by lung particle uptake in %ID/g. (f) *Ex vivo* quantification of lung, liver, and spleen luminescence for mice treated with 5μg pDNA at 1d (left) and 14d (right), respectively, shows strong retention of lung luminescence over time with marked decreases in liver and spleen signal over the same period. Both IgG and bare DNA-LNP luminescence decreased to ∼2x background by day 3 and were thus excluded at day 14. For panel (d), an unpaired *t*-test with Welch’s correction was performed, and lines are drawn through median. In panel (e), a one-way ANOVA with Tukey’s multiple comparisons test was performed. In panel (f), a two-way ANOVA with Tukey’s multiple comparisons test was performed. All data include n = 3 and represent mean ± SEM; * = *p* < 0.05, ** = *p* < 0.01, *** = *p* < 0.001, **** = *p* < 0.0001.

To confirm that our observed luminescence was primarily in the lungs, we IV treated mice with 5μg of pDNA loaded into PECAM-targeted or untargeted IgG DNA-LNPs. At either 1 or 14 days after treatment, mice were euthanized to measure the total flux of *ex vivo* luminescence of individual organs via IVIS (quantification method provided in **Figure S6**). At both days 1 and 14, the majority of luminescence emitted from the lungs, with a drop in luminescence from ∼8 × 10^5^ photons s^-1^ at day 1 to ∼5 × 10^5^ photons s^-1^ at day 14 (**Figures 2b, 2f**). This was significantly higher than the lung expression measured from bare and IgG DNA-LNPs and untreated control mice, both of which emitted luminescence ∼10^4^ photons s^-1^ at 1d. Given the sharp decline of total *in vivo* bare and IgG DNA-LNP luminescence to a mere ∼2x background by day 3, *ex vivo* quantifications at day 14 were not performed.

Surprisingly, our *ex vivo* quantifications revealed considerable decrease in liver luminescence in mice treated with αPECAM DNA-LNPs over the 14-day period, dropping a full order of magnitude by day 14. This decrease in liver signal is significantly larger than the concurrent decrease in lung signal. Thus, the specificity of our αPECAM DNA-LNP lung expression increases over time, with a nearly 5-fold increase in the quantified lung:liver expression ratio over 14 days (**Figure 2d**). Notably, αPECAM DNA-LNPs also have a significantly higher lung:liver expression ratio than αPECAM mRNA-LNPs at peak expression^27^. When normalized by the ID/g delivery to the lungs, this difference becomes even more pronounced, demonstrating the increased specificity and efficiency of our DNA delivery platform. (**Figure 2e**).

### 2.3 Targeting VCAM-1 redirects the delivery and expression of DNA-LNPs to the spleen and brain

In order to show the generalizability of our platform, we swapped the targeting antibody from αPECAM to anti-VCAM-1 (αVCAM) mAbs. VCAM-1 is a cell adhesion molecule, similar to PECAM-1, whose expression is highly upregulated during pathological conditions such as ischemic stroke, intracerebral hemorrhage, or Alzheimer’s and Parkinson’s Disease^30,32,35,36^. Previous studies have shown that drug delivery systems targeting VCAM-1 accumulate in the spleen^30,37^. However, αVCAM nanocarriers also experience a significant increase in brain uptake compared to controls^29,31,32^. Our results similarly show that radiolabeled, IV-administered αVCAM DNA-LNPs (50 mAbs per particle) accumulate in the spleen of naive mice 30 minutes after injection at ∼85 %ID/g (**Figure 3a**). This was significantly more than αPECAM and untargeted IgG DNA-LNPs, which both have a %ID/g of ∼50% in the spleen. αVCAM DNA-LNPs also exhibited over threefold higher brain targeting compared to both αPECAM and untargeted IgG nanoparticles (**Figure 3d**).

**Figure 3.**
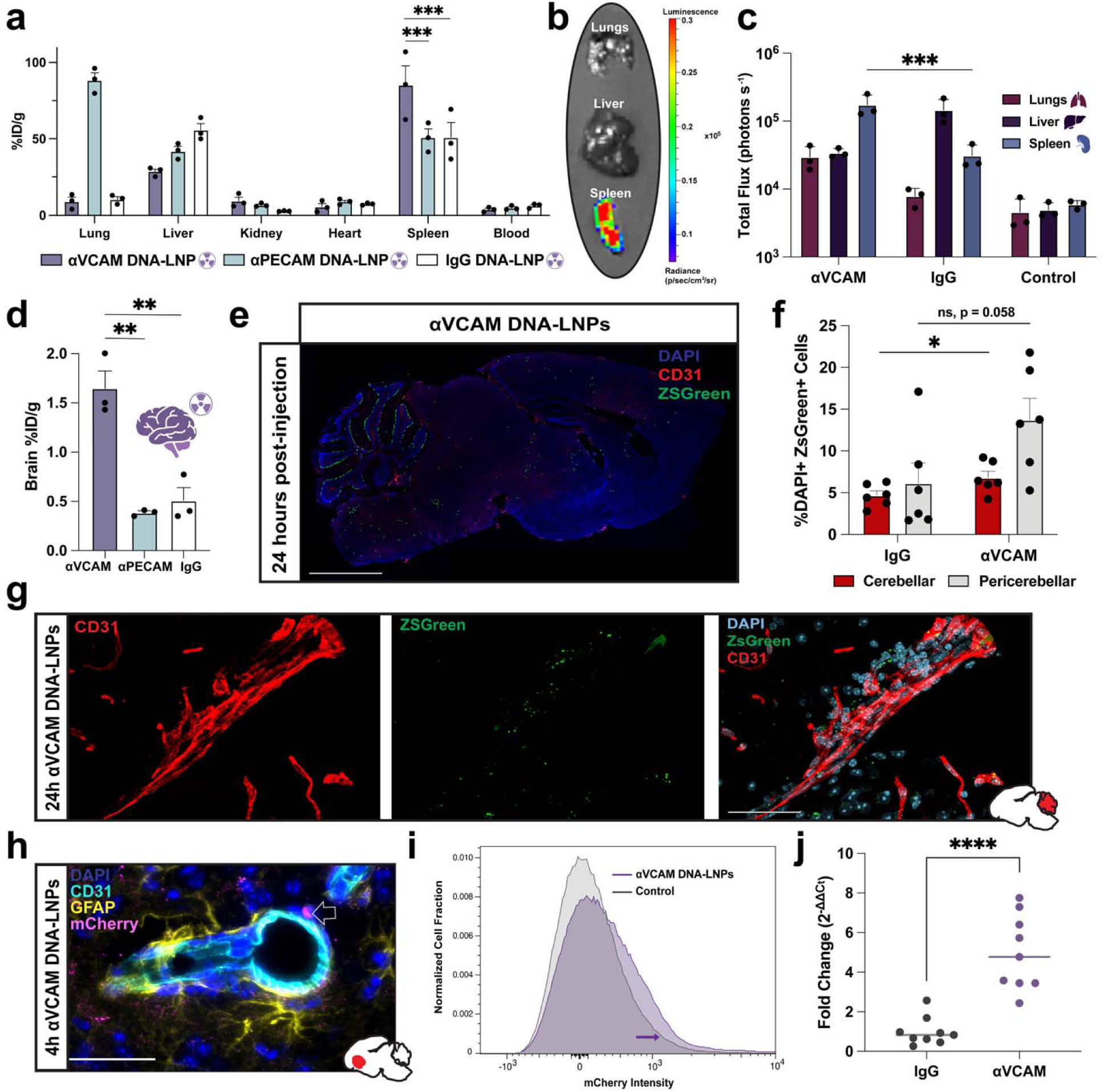
VCAM targeted DNA-LNPs induce elevated transgene expression in both the spleen and brain. (a) Biodistribution of αVCAM DNA-LNPs, αPECAM DNA-LNPs, and IgG DNA-LNPs via ^125^I radiotracing in naive BALB/c mice reveals considerable shifts in particle distribution based on targeting moiety. αVCAM DNA-LNPs exhibit the highest spleen uptake compared to other particles. (b) *Ex vivo* bioluminescence imaging 1d post-injection with αVCAM DNA-LNPs containing 5μg luciferase pDNA shows the highest expression in the spleen. (c) Quantification of *ex vivo* images underscores differences in major organs of mice treated with αVCAM versus untargeted IgG DNA-LNPs, with αVCAM DNA-LNPs exhibiting significantly higher expression in the spleen compared to both the lungs and liver. Naive, luciferin-treated controls are shown for comparison. (d) Radiotracing analysis in the brain reveals a >3x increase in particle uptake of αVCAM DNA-LNPs compared with both αPECAM and IgG DNA-LNPs. (e) Immunofluorescent (IF) sagittal brain section (20X resolution) 24 hours after treatment with αVCAM DNA-LNPs loaded with pDNA encoding Cre recombinase. Merge depicts DAPI (blue), ZsGreen dot render (green), PECAM-1 (red) (scale bar, 1mm). (f) Quantification comparing (peri)cerebellar expression of Cre-induced ZsGreen in αVCAM and IgG DNA-LNPs. (g) 60X images highlighting endothelial cells (CD31) and ZsGreen signal demonstrating colocalization of ZsGreen expression in endothelial cells and surrounding neurovasculature (scale bar, 50µm). (h) 40X IF image from the brain of a naive BALB/c mouse 4 hours after treatment with αVCAM DNA-LNPs loaded with mCherry pDNA. Merge shows DAPI (blue), CD31 (cyan), mCherry (magenta), and astrocyte marker GFAP (yellow) (scale bar, 50µm). mCherry expression is apparent in the perivascular space; arrow indicates mCherry expression. (i) Comparison of total mCherry fluorescence in the brain cells of mice that are either naive (control, grey) or treated with 25μg αVCAM mCherry DNA-LNPs (purple) measured via flow cytometry, illustrating a shift in mCherry expression in αVCAM DNA-LNP treated brains. (j) Reverse Transcription quantitative Polymerase Chain Reaction (RT-qPCR) data of mCherry mRNA levels in the brains of mice treated with 25μg of either αVCAM mCherry DNA-LNPs or IgG DNA-LNPs. αVCAM DNA-LNP treated brains exhibit a ∼5-fold increase in mCherry mRNA levels compared to untargeted controls. For panels (a) and (c), two-way ANOVAs with Tukey’s multiple comparisons test were performed. In panel (d), a one-way ANOVA with Tukey’s multiple comparisons test was performed. In panel (f), an unpaired *t*-test with Welch’s correction was performed, and lines are drawn through median. In panel (j), an unpaired *t*-test was performed on ΔCt, where ΔCt = log_2_(fold change), and lines are drawn through median. All data include n = 3 biological replicates, except panel (f), which includes n = 2 biological replicates; panels (f) and (j) contain two and three technical replicates per biological replicate, respectively. All data represent mean ± SEM; ** = *p* < 0.01, *** = *p* < 0.001, **** = *p* < 0.0001.

To quantify the expression of VCAM-targeted nanoparticles, we again IV-treated mice with 5μg of luciferase pDNA loaded into αVCAM DNA-LNPs and measured *ex vivo* luminescence of individual organs via IVIS 1 day following treatment. At this time and dose, we only observed elevated luminescence in the spleen at ∼2 × 10^5^ photons s^-1^, which was significantly higher than mice treated with 5μg of luciferase pDNA loaded into untargeted IgG DNA-LNPs and naive, luciferin-treated control mice (**Figure 3b-c**). We did not detect luminescence in the brain, which is expected given (1) the relatively low dose of pDNA and (2) the detection sensitivity of IVIS for organs with lower nanoparticle uptake.

To show that our αVCAM DNA-LNPs express in the brain, we used immunofluorescence (IF). Ai6 mice, which express an enhanced green fluorescent protein (ZsGreen) following Cre-mediated recombination of a STOP cassette, were intravenously injected with αVCAM or untargeted IgG DNA-LNPs loaded with 5μg of pDNA encoding Cre recombinase. 24 hours after treatment, brains were harvested and stained with DAPI and fluorescent antibodies against PECAM-1 (CD31), and a representative sagittal dot render was produced (**Figure 3e**). Based on the areas of highest apparent expression, we imaged multiple randomly selected fields at 20X magnification in cerebellar and pericerebellar regions prior to running a custom macro that calculated the percent of DAPI+ cells that were also positive for ZsGreen. Our quantification shows significantly increased cerebellar ZsGreen expression in the brains of mice treated with αVCAM DNA-LNPs compared to IgG DNA-LNPs (**Figure 3f**). A representative confocal image of a brain blood vessel from a mouse treated with αVCAM DNA-LNPs shows expression of ZsGreen either colocalized with or adjacent to endothelial cells (**Figure 3g**). In contrast, control brains from IgG DNA-LNP–treated mice did not show comparable endothelial expression (**Figure S8**).

Similarly, we repeated IF using αVCAM or untargeted IgG DNA-LNPs loaded with mCherry pDNA, encoding a red monomeric fluorescent protein, 4 hours post-injection in naive BALB/c mice. This alternate approach also shows expression observed within the neurovasculature, which is significant as mCherry is a true transgene as opposed to the constitutive expression of Cre-induced ZsGreen in Ai6 mice (**Figure 3h, Figure S7**). These images confirm that αVCAM DNA-LNPs target and express in different regions of the brain endothelium as early as 4 hours post-treatment. Quantifications for immunofluorescence comparisons between αVCAM and IgG mCherry DNA-LNPs trend significance, likely due to the earlier time point and lower sensitivity of mCherry (**Figure S8-9**). To provide an additional line of evidence of brain expression, we utilized flow cytometry to detect mCherry fluorescence in single-cell suspensions of the brains of either naive (control) mice or mice treated with 25μg αVCAM mCherry DNA-LNPs 24 hours prior. The brain cells of αVCAM DNA-LNP treated mice have a noticeable shift in mCherry fluorescence compared to controls, indicating low but detectable transgene expression (**Figure 3i**). To more sensitively assess mCherry expression, we next performed Reverse Transcription quantitative Polymerase Chain Reaction (RT-qPCR) on brain homogenates from naive and αVCAM mCherry DNA-LNP treated mice 24 hours post-injection (**Figure 3j**). αVCAM DNA-LNP treated brains had ∼5-fold increase in mCherry mRNA levels compared to untargeted IgG DNA-LNPs, confirming substantial, specific DNA delivery and transcription consistent with the expression patterns observed by IF and flow cytometry.

Taken together, these findings highlight the robustness of antibody targeting, and the potential of αVCAM DNA-LNPs to serve as a drug delivery system, notably to treat chronic pathologies of the brain.

### 2.4 Fabs improve the delivery, expression, and specificity of lung targeted DNA-LNPs

We repeated prior experiments with αPECAM Fabs conjugated to our DNA-LNPs rather than mAbs. Fabs are the antigen-binding fragments of full-length antibodies, and therefore are smaller than mAbs (50 kDa vs 150 kDa) and lack a fragment crystallizable (Fc) region, which is the portion of the antibody that interacts with cell surface receptors on immune and other cell types^38–40^ **(Figure 4a)**. We hypothesized that Fabs would improve the specificity of αPECAM DNA-LNPs by eliminating Fc-mediated interactions of the LNPs with targets other than PECAM. αPECAM Fabs conjugated to DNA-LNPs as well as mAbs, with a conjugation efficiency of >80% (**Figure S1**). Fabs were added to DNA-LNPs at a concentration of 100 Fabs per particle, in order to match the number of binding domains on the LNPs with 50 mAbs/particle tested above (**Figure 4a)**. Radiolabeled αPECAM-Fab DNA-LNPs targeted the lungs of naive mice at ∼80% ID/g 30 minutes after IV treatment, matching mAb-conjugated nanoparticles (**Figure 4b**).

**Figure 4.**
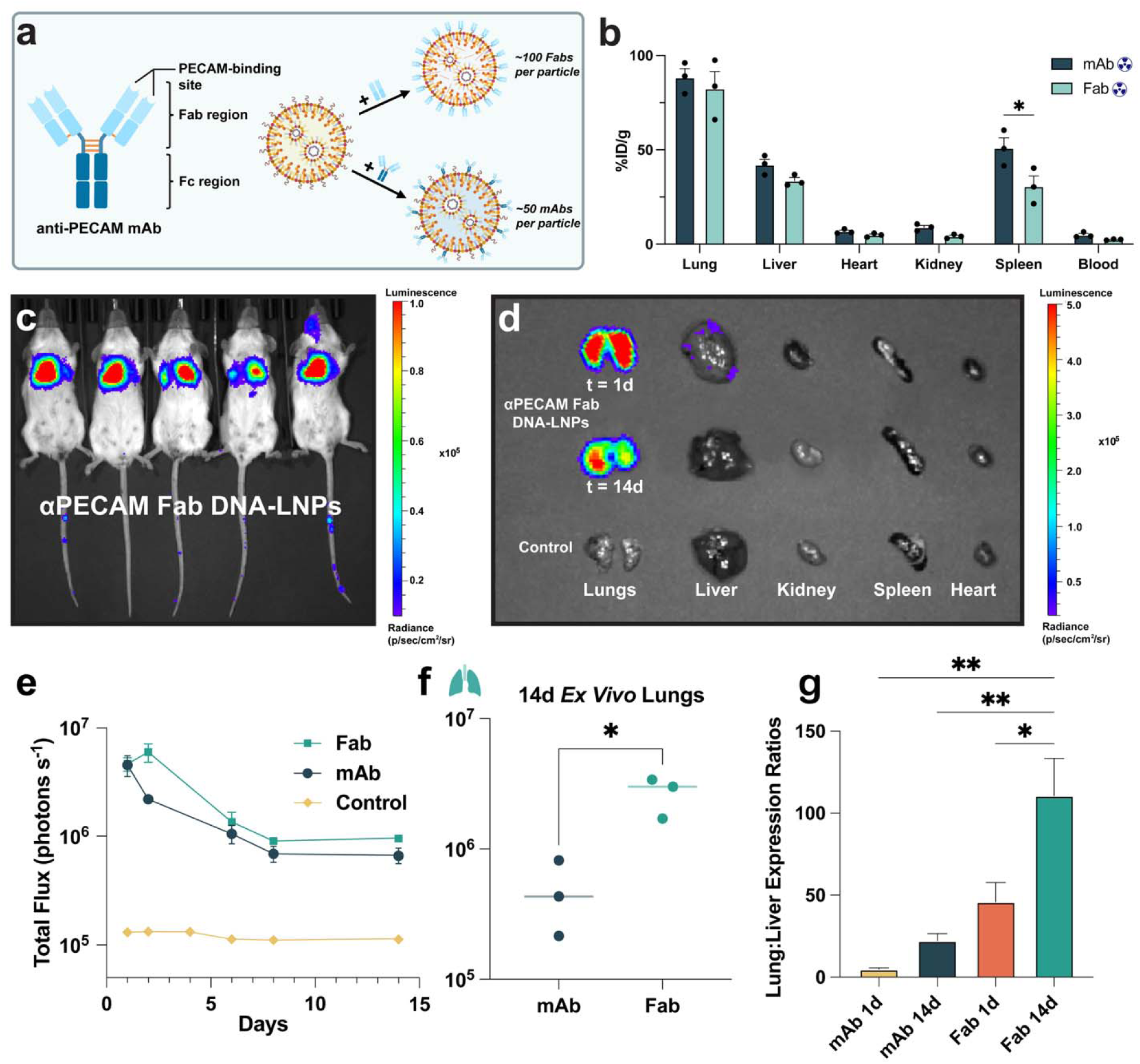
Conjugating anti-PECAM Fabs to DNA-LNPs immensely increases both lung specificity and expression. (a) Schematic illustrating the structure and conjugation of αPECAM mAbs vs Fabs to DNA-LNPs. Notably, Fabs are ∼50 kDa compared to mAbs (∼150 kDa), and lack the fragment crystallizable (Fc) region of full-length antibodies. (b) Biodistribution via radiotracing of ^125^I-labeled Fab and mAb αPECAM DNA-LNPs reveals similar lung accumulation by both particles, and decreased uptake of Fab DNA-LNPs in the spleen. (c) Representative IVIS image of BALB/c mice treated with 5μg of luciferase pDNA loaded into αPECAM Fab DNA-LNPs shows strong luminescent signal in the thoracic cavity, especially in the lungs. (d) Representative *ex vivo* image displaying lung luminescence from αPECAM Fab DNA-LNP-treated mice that persists over time. (e) Comparison of *in vivo* luminescence shows superiority of αPECAM Fab DNA-LNPs compared to mAb DNA-LNPs. (f) *Ex vivo* quantification confirms superior lung expression at 14d in Fab condition, showing roughly an order of magnitude increase in luminescence compared with mAb DNA-LNPs at the same time point. (g) Lung-to-liver expression ratios quantified at 1d and 14d via *ex vivo* luminescence show increasing lung specificity by αPECAM Fab DNA-LNPs over time, and overall higher ratios compared to αPECAM mAb DNA-LNPs. In panel (b), a two-way ANOVA with Tukey’s multiple comparisons test was performed. For panel (f), an unpaired *t*-test with Welch’s correction was performed, and lines are drawn through median. In panel (g), a one-way ANOVA with Tukey’s multiple comparisons test was performed. All data include n = 3 and represent mean ± SEM; * = *p* < 0.05, ** = *p* < 0.01.

Once again, we used IVIS to investigate *in vivo* transgene expression of αPECAM-Fab DNA-LNPs loaded with 5μg of luciferase pDNA. One day after IV injection in naive mice, luminescence was concentrated in the thoracic cavity, with the majority of the signal emitting from the lungs, visibly outlining the organ (**Figure 4c**). This is confirmed by *ex vivo* analysis of individual organs 1 day post-treatment, which demonstrates that the majority of signal originates from the lungs, with low-level luminescence in the liver and spleen (**Figure 4d**). The total flux of the overall *in vivo* luminescence in αPECAM-Fab DNA-LNP treated mice after one day was very similar to mAbs at ∼5 × 10^6^ photons s^-1^, although Fab expression peaked higher after 2 days at ∼7 × 10^6^ photons s^-1^ (**Figure 4e**). The duration of αPECAM-Fab DNA-LNP luminescence followed a nearly identical trend to mAbs, steadily declining until day 7 before plateauing at 10^6^ photons s^-1^ from days 7-14.

Notable differences between Fab- and mAb-conjugated DNA-LNPs became more apparent when analyzing the transgene expression of *ex vivo* organs over time. 14 days post-injection, the lung luminescence in mice treated with αPECAM-Fab DNA-LNPs was nearly an order of magnitude higher than mAbs (**Figure 4f**). Moreover, the specificity of αPECAM-Fab DNA-LNP lung expression was substantially higher than that of mAbs, with their lung:liver expression ratio increasing from 27 at day 1 to 111 at day 14, compared to 5 at day 1 and 22 at day 14 for mAbs (**Figure 4g**).

To further visualize the expression of our αPECAM-Fab DNA-LNPs in the lungs, we performed immunofluorescence. Mice were IV-injected with αPECAM-Fab DNA-LNPs loaded with 5μg of pDNA encoding mCherry. Sections prepared from lungs 4 hours after treatment showed expression of mCherry in endothelial cells of the pulmonary vasculature (**Figure S7**). In totality, these findings emphasize that Fabs are the more specific, higher expressing, and overall superior moiety for lung-targeted DNA-LNPs.

### 2.5 PECAM-targeted Fab DNA-LNPs primarily deliver to and express in pulmonary endothelial cells

To probe cell specificity of our optimized targeting formulation with αPECAM Fabs, we utilized flow cytometry to identify the cell types that our αPECAM-Fab DNA-LNPs are delivered to and ultimately expressed in. First, we formulated αPECAM-Fab DNA-LNPs with Alexa Fluor 488 (AF488) fluorescent lipid to track the cellular distribution and uptake of the nanoparticle. These LNPs were IV-injected into naive mice at a 5μg pDNA dose, and 30 minutes later the animals were sacrificed and perfused to prepare the lungs and liver into single cell suspensions. Antibody stains identified general immune cells (anti-CD45), monocytes and macrophages (anti-CD64), neutrophils (anti-Ly6G), endothelial cells (anti-CD31), and epithelial cells (anti-EpCAM) to determine which cells αPECAM-Fab DNA-LNPs were delivered to (**Figure S11**). Out of all the LNP+ cells in the lungs, 77% were endothelial, 13% were CD45+/CD64-/Ly6G-immune cells (labeled, “other immune cells”), and 3% were neutrophils (**Figure 5a, Left**). In the liver, LNPs were primarily delivered to CD45-/CD31-/EpCAM-cells (labeled, “other cells”) at 53%, followed by endothelial cells at 29%. The “other cells” in the liver are predominantly hepatocytes. This analysis illustrates that αPECAM-Fab DNA-LNPs are highly endothelial-specific, especially in the lungs.

**Figure 5.**
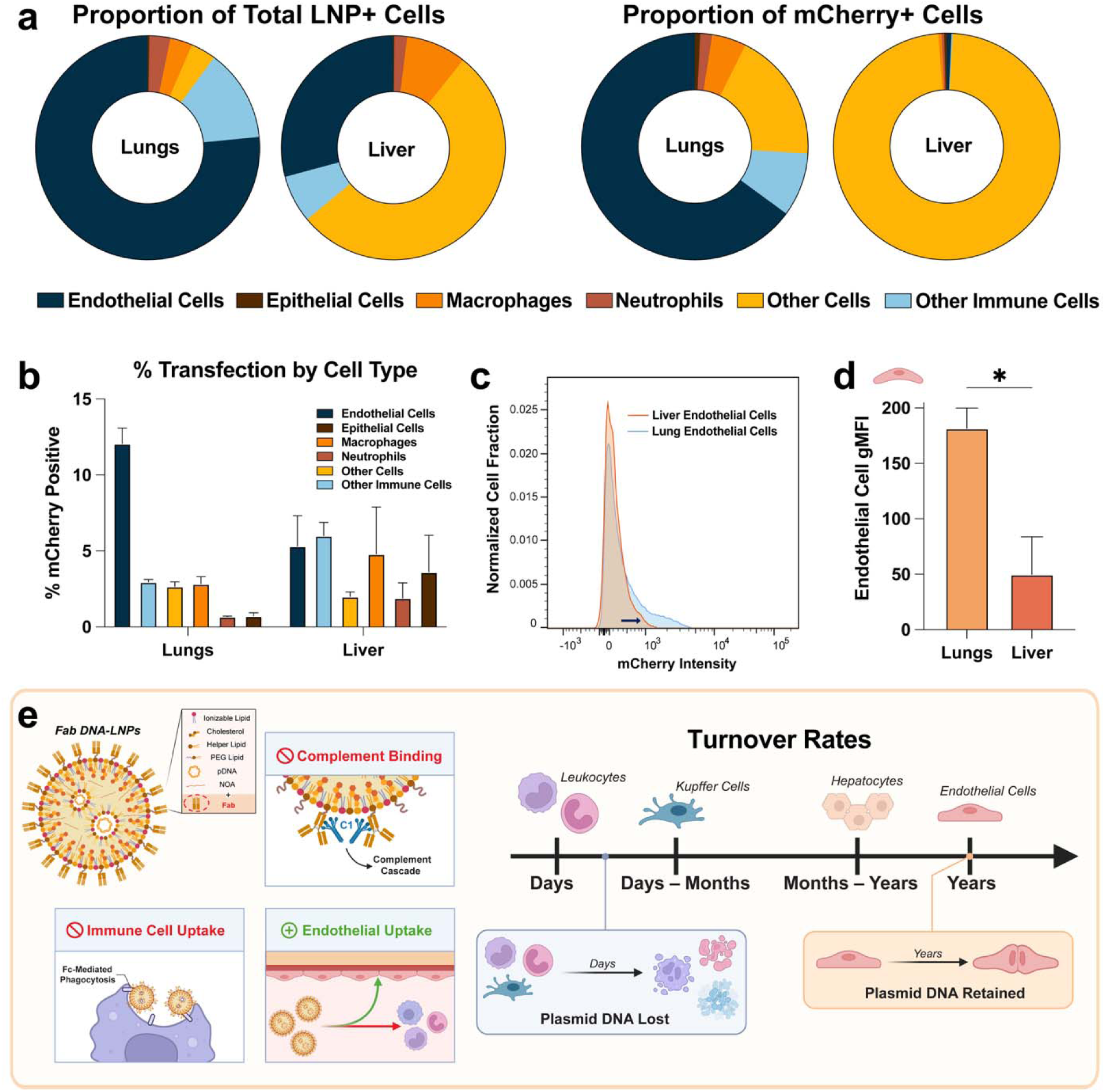
Anti-PECAM Fabs facilitate uptake and transgene expression of DNA-LNPs with specificity to lung endothelial cells. (a) Flow cytometry analysis tracking the (left) proportion of cells positive for fluorescent αPECAM-Fab DNA-LNPs and (right) proportion of cells positive for mCherry pDNA expression induced by αPECAM-Fab DNA-LNPs. Left panel: >75% of LNP+ cells in the lungs are endothelial (CD45-/CD31+), while the majority of LNP+ cells in the liver are “other cells” (CD45-/CD31-/EpCAM-). Right panel: mCherry pDNA transgene expression by cell type reveals that the majority of mCherry-expressing cells are endothelial in the lungs and “other cells” in the liver. Note that hepatocytes in the liver were not specifically stained for and thus fall under the “other cells” population. (b) Alternative view of the data presented in (a, right panel) showing the percentage of mCherry+ cells in each cell type in the lungs and liver, revealing the highest fraction of mCherry positivity to be in lung endothelial cells (ECs). (c) Comparison of total mCherry fluorescence in the ECs of the liver and lungs, illustrating a significant shift in mCherry expression in lung ECs. (d) Quantification of mCherry fluorescence in lung and liver endothelial cells by geometric mean fluorescence intensity (gMFI), demonstrating significantly higher expression in lung ECs compared to liver ECs. (e) Graphic illustrating the leading hypothesis for superior lung specificity of αPECAM-Fab DNA-LNPs and increasing lung specificity over time. Left: Fabs, which lack the Fc domain of full-length mAbs, avoid classical pathway complement activation and engagement with Fc receptors on immune cells, resulting in superior lung EC binding and expression compared to mAbs^41–43^. Right: Lung ECs maintain persistent transgene expression over time due to their slower turnover rate, while more short-lived cells in the liver lose plasmid DNA and thus expression^44–46^. Additionally, turnover mechanisms affect whether pDNA is destroyed, as in apoptosis, or diluted, as in cell division. For panel (d), an unpaired *t*-test with Welch’s correction was performed. All data include n = 3 and represent mean ± SEM; * = *p* < 0.05.

Because LNP delivery does not always lead to transgene expression, we also utilized flow cytometry to detect the expression of αPECAM-Fab DNA-LNPs in the same cell populations. 5μg of mCherry pDNA were loaded into αPECAM-Fab DNA-LNPs and IV administered into naive mice. 1 day after treatment, the animals were sacrificed and perfused to prepare single cell suspensions for flow cytometry using the same parameters as before. 65% of mCherry expressing cells in the lungs were endothelial, followed by “other cells” at 19%. (**Figure 5a, Right**). In totality, 12% of all pulmonary endothelial cells recovered expressed mCherry, which is higher than any other cell population (**Figure 5b**). In the liver, nearly all of the mCherry expressing cells were from the “other cells” population, at 98% (**Figure 5a, Right**). This reveals an interesting discrepancy, as αPECAM-Fab DNA-LNPs delivered relatively well to ECs in the liver yet expressed poorly. While surprising, this phenomenon could be explained by differences in the endothelial cell expression phenotype between the liver and lungs. Our flow data supports this hypothesis, as our recovered population of pulmonary endothelial cells have a noticeable shift in mCherry expression compared to liver endothelial cells, and also have a significantly higher geometric mean fluorescence intensity (gMFI) (**Figures 5c-d).**

Overall, our flow data uncover a theory explaining why the lung specificity of αPECAM-Fab DNA-LNPs is superior to that of mAbs and increases over time. Fabs can evade recognition by the innate immune system due to their lack of an Fc domain^41,42^ (**Figure 5e**). Therefore, αPECAM-Fab DNA-LNPs are less likely to be taken up by phagocytes expressing Fc receptors and subsequently sequestered to reservoirs of immune cells in the liver^43^. Instead, they exhibit superior binding and uptake by pulmonary endothelial cells, which may account for the greater lung specificity of Fab-conjugated nanoparticles relative to mAbs. This hypothesis also explains why lung specificity increases over time in general, as the half-lives of immune cells and other cells are significantly shorter than endothelial cells, which remain quiescent for years^44,45^ (**Figure 5e**). Therefore, the expression of cells in the liver diminishes over the course of 14 days, while pulmonary endothelial cell expression persists^46^. Furthermore, our DNA plasmids contain the cytomegalovirus (CMV) promoter, which has been reported to be rapidly silenced in liver hepatocytes when delivered via viral vectors^47^. This means the effect of our αPECAM-Fab DNA-LNPs in the lungs will persist, driven by high-expressing, long-lived endothelial cells, despite its reduction in the liver and other organs.

## 3. Discussion

While mRNA-LNP therapies have seen success for a few applications where transgene expression is meant to be short-lived, the hours-long half-life of mRNA limits clinical use in the treatment of chronic diseases, especially those without a clear genetic target. Furthermore, regardless of cargo, LNPs primarily accumulate in the liver, limiting their therapeutic relevance in extrahepatic diseases. Thus, following our prior work mitigating the morbidity and mortality of DNA-LNPs by loading them with the STING inhibitor NOA, we sought to extend this platform by targeting our novel drug delivery system to various vascular beds, focusing primarily on the lungs. In doing so, we show effective, organ-specific transgene expression that underscores the potential for targeted DNA-LNP therapies in chronic vascular diseases of multiple organs.

Using bioluminescence, we demonstrate that our αPECAM DNA-LNPs induce high transgene expression in the lungs that persists for weeks, with significant increases in lung-versus-liver expression over time. Surprisingly, the target specificity of our αPECAM DNA-LNPs is markedly greater than that of our prior αPECAM mRNA-LNPs. This difference is amplified with time, suggesting that DNA’s chronicity imparts specificity whereas mRNA is simply degraded. Single-cell analysis confirmed that the vast majority of DNA expression in the lungs was confined to target endothelial cells, which are more quiescent than other transfected cell types^44,45^. From these data, we can extract two main points: (1) PECAM targeting is a highly effective method to achieve lung delivery of DNA-LNPs, and (2) targeting DNA-LNPs to the vascular endothelium prolongs transgene expression by confining DNA delivery to cells that will not actively apoptose or divide. The corollary to this second point is that any organ whose endothelium is effectively targeted will express a transgene with higher specificity over time, as the DNA is either destroyed or diluted in other cell types that have shorter half-lives or undergo frequent divisions. These phenomena are not applicable to mRNA given that its half-life is shorter than most cell types. Thus, the endothelial cell is an optimal target for delivery of DNA-LNPs.

We expanded on our finding by targeting the same PECAM-1 epitope with Fabs in place of mAbs. We hypothesized that removing the Fc region from mAbs would limit phagocytic uptake and complement opsonization of our LNPs^38–40^. Indeed, using Fabs improved both lung specificity and expression of our DNA-LNPs. We attribute this phenomenon to the increased endothelial targeting of αPECAM-Fab DNA-LNPs, as they are less visible to the body’s innate immune mechanisms. Interestingly, the geometric mean fluorescent intensity of lung endothelial cells is more than triple that of liver endothelial cells, indicating fundamental differences in how the endothelial cells of various organs process DNA delivered by LNPs. Given the unique role that liver sinusoidal endothelial cells (LSECs) play in the clearance of blood-borne waste, they exhibit a highly endocytic and degradative phenotype^48,49^. In fact, LSECs have previously been identified as the primary cell type by which naked plasmid DNA is cleared in rodents^50^. This offers an explanation as to why LSECs avidly take up αPECAM DNA-LNPs but do not express their DNA cargo to the same extent as lung endothelial cells. These differences amongst endothelial cells are not explored any further herein and will need to be studied in future work. Likewise, the effect that the CMV promoter has on the inhibition of expression in hepatocytes warrants further investigation^51^. Since cytomegalovirus primarily infects hepatocytes, it is possible that these cells have evolutionarily adopted rapid silencing mechanisms of the CMV promoter. Thus, we believe the higher, persistent transgene expression exhibited in pulmonary endothelial cells is a combination of these, among numerous other factors.

Finally, we aimed to show the flexibility of our platform by swapping out the PECAM-1 targeting moiety with one against VCAM-1. αVCAM DNA-LNPs showed highly specific spleen expression and increased brain uptake and expression compared with αPECAM and IgG DNA-LNPs. Although not shown here, VCAM-1 is heavily upregulated in the brain microvasculature during inflammatory states, which can greatly increase nanoparticle uptake^28–32^. When considering that chronic diseases are generally accompanied by global low-level inflammation, it is conceivable that VCAM-targeting would improve brain specificity and expression in chronic neurological disease models, warranting future use of our αVCAM DNA-LNPs in studies of brain diseases^52^.

While this study achieves specific endothelial targeting of DNA-LNPs, additional work is needed to expand on our platform technology. Further studies distinguishing the various subtypes of endothelial cells transfected (*e.g.*, arterial, capillary, venous) will determine the pathologies targeted in future preclinical work. For example, if pulmonary arterial endothelial cells are significantly transfected, we can adapt our platform to deliver therapeutics in disease models of pulmonary hypertension. Likewise, while our targeted DNA-LNPs target the organ of interest with high endothelial transfection, promoters could further enhance endothelial specificity—an advantage enabled by the inherent flexibility of DNA cargo. Similarly, we can further tailor the delivery of our DNA-LNPs to sites of inflammation in disease models by choosing the appropriate ligand. Previous research from our lab demonstrated that PECAM-targeted liposomes achieve broader pulmonary uptake whereas ICAM-targeted liposomes preferentially localize to inflamed tissue^53^. Therefore, it will be important to explore whether similar targeting dynamics apply to DNA-LNPs in inflammatory disease models. We envision that combining our targeted DNA-LNPs with advanced DNA and ligand engineering will enable us to leverage endothelial cells as biofactories that secrete therapeutics directly onto parenchymal cells, specifically at the site of inflammation or disease.

Collectively, our study underscores key advantages of DNA-LNP delivery over mRNA and greatly expands the therapeutic prospects of DNA-LNPs to include chronic vascular diseases of the lungs, spleen, and brain. Furthermore, we have shown that simply swapping targeting moieties is an effective tool to alter the biodistribution of DNA-LNPs and induce organ-specific transgene expression that invariably increases over time when targeting endothelial cells.

## 4. Experimental Section

### Antibodies

Anti-mouse-PECAM-1/CD31 (clone 390) monoclonal antibodies and fragment antigen-binding regions were obtained from Sino Biological. Control rat IgG was purchased from Invitrogen. Anti-mouse-VCAM-1 (clone MK2.7) was produced by culturing hybridoma cells, purified using protein G sepharose (GE Healthcare Bio-Sciences, Pittsburgh, PA) and dialyzed in PBS.

### Antibody modification

Antibodies (mAbs and Fabs) were functionalized with DBCO (Dibenzocyclooctyne) for conjugation to LNPs by mixing the antibody with a 7-fold molar excess of DBCO-PEG4-NHS Ester (BroadPharm) and a 0.5 molar excess of Alexa Fluor 594 NHS Ester (Thermo Fisher) for 1 hour at room temperature with rotation. Unreacted DBCO and fluorophore was removed by washing three times with 10x excess PBS in 10 kDa molecular weight cut off Amicon Ultra centrifugal filters (MilliporeSigma).

### Preparation of mRNA/DNA-LNPs

Lipids (Echelon Biosciences for ionizable lipids, Avanti Polar Lipids for others), dissolved in ethanol, were combined in the molar percentages described in **Figure 1A or Figure S2**. For DNA LNPs, 9(10)-nitrooleic acid (NOA) (Echelon Biosciences), dissolved in ethanol, was added to the lipid mixture at a drug-to-total lipid ratio of 0.2 (mole-to-mole). Messenger RNA (TriLink) or plasmid DNA (Aldevron) was dissolved in buffer (50 mM citrate buffer, pH 4). LNPs were formulated using microfluidics (NanoAssemblr Ignite, Precision Nanosystems) at a total flow rate of 6 mL/min, a flow rate ratio of 1:3 (lipid:nucleic acid mixture), and total lipid: nucleic acid ratio of 40-to-1 (wt/wt). Following formation, LNPs were dialyzed with 1x PBS in a 10 kDa molecular weight cut-off cassette (Life Technologies) for 2 hours.

### LNP Characterization

Following formulation, nanoparticle size and polydispersity index (PDI) was measured via dynamic light scattering (DLS) using the Zetasizer Nano ZS (Malvern Instruments Ltd). The encapsulation efficiency and concentrations of nucleic acid within the LNPs were determined using either Quant-iT PicoGreen dsDNA (for DNA LNPs) or Ribogreen (for mRNA) assays (Invitrogen).

### Antibody Radiolabeling

Rat IgG isotype control antibodies (Invitrogen) were radiolabeled with Na^125^I using Pierce’s Iodogen radiolabeling method. To summarize, tubes were coated with 100μg Iodogen. IgG (at a concentration of between 1–2 mg/mL) and Na^125^I (0.25μCi/μg protein) were incubated for 5 mins on ice. Unreacted materials were purified using 7 kDa Zeba desalting columns (Thermo Fisher Scientific). Thin-layer chromatography was used to confirm that all antibodies had >90% radiochemical purity prior to use.

### Antibody Conjugation

Before conjugation, the particle concentration of azide-functionalized LNPs were measured via nanoparticle tracking analysis (NTA) using a Nanosight NS300 (Malvern Panalytical). LNPs were then incubated with DBCO-modified antibodies at the appropriate Ab:LNP ratio overnight at 4°C. Unbound antibodies were purified from the conjugation by passing the mixture through a size exclusion column loaded with Sepharose CL-4B (Cytiva), followed by 24 consecutive 1 mL elutions with PBS that were individually collected. Conjugation efficiency was quantitatively assessed by measuring the ratio of the area under the curve for the fluorescent or radiolabeled antibody in the LNP fractions (6–8 mL) to the signal in the entire 24 mL elution. The conjugation efficiency for targeted LNPs range between 70-95%, depending on LNP and antibody batch, resulting in approximately 50 mAb/LNP or 100 Fabs/LNP.

### Animals

All animal experiments strictly adhered to the guidelines established in the Guide for the Care and Use of Laboratory Animals (National Institutes of Health). Euthanasia techniques will strictly follow the AVMA Guidelines for the Euthanasia of Animals: 2020 Edition. Approval for all animal procedures was obtained from the University of Pennsylvania Institutional Animal Care and Use Committee. Female Naive BALB/c and B6.Cg-Gt(ROSA)26Sortm6(CAG-ZsGreen1)Hze (Ai6 for short) mice aged 8–12 weeks weighing 18– 25g, were procured from The Jackson Laboratory for all studies. The mice were housed in a controlled environment run by the University Laboratory Animal Resources (ULAR) facility and were maintained at temperatures between 22 °C and 26 °C with a 12-h light/dark cycle, and provided with ample access to food and water.

### In Vivo Studies

For *in vivo* studies, although our LNPs do not produce unacceptably high levels of distress, the following guidelines were strictly adhered to by euthanizing all animals that met any of the following criteria: 1. Animals have a body condition score of 1; 2. Animals have a body condition score of 2 in addition to other signs of distress such as hunched posture, porphyrin staining, inactivity, ruffled hair coat, or dehydration; 3. Animals show overt signs of injury (redness, swelling); 4. Animals show weight loss >= 20%. All intravenous (IV) injections were done retro-orbitally by injecting into the retro-bulbar sinus. All euthanizations were performed using both cervical dislocation and exsanguination by cutting the IVC and descending aorta.

### Biodistribution Studies

For biodistribution studies, Na^125^I IgG was conjugated to LNPs at a ratio of 5 IgG/LNP as a signal to track particle distribution, alongside the targeted or control antibody at the usual 50 mAb/LNP or 100 Fab/LNP. Following Antibody-LNP conjugation and purification, mice were IV injected with LNPs at a dose of approximately 2.5μg nucleic acid. Thirty minutes after injection, animals were sacrificed. Blood was collected in EDTA-coated tubes (Thermo Fisher), and organs (lung, liver, spleen, heart, kidney, and brain) were collected and weighed. Tissue distribution of injected materials was determined by measuring the radioactivity in the blood and organs using a Wizard 2470 Gamma Counter (PerkinElmer). Organ uptake was calculated as percent injected dose normalized to the mass of tissue (%ID/g tissue).

### In Vivo Imaging System (IVIS) for Luciferase Expression

Prior to IVIS imaging, mice were intravenously injected with targeted or control DNA-LNPs loaded with pDNA encoding for Luciferase (Aldevron) at a dose of 2.5 or 5μg. 1-14 days post treatment, mice were put under using 3% isoflurane-induced and intraperitoneally injected with 100µl of 30 mg/mL D-luciferin sodium salt (Regis Technologies). Anesthetized mice were placed in an IVIS Spectrum machine (Revvity) face up (to view the thorax) and imaged for chemiluminescence in the target area every minute with automatically determined exposure time for 10–14 images, until the signal reached the peak intensity. Revvity LivingImage software (version 4.8.2) was used to analyze images.

### Ex Vivo Imaging

Following IVIS imaging on either day 1 or 14, mice that were already intraperitoneally injected with 100µl of 30 mg/mL D-luciferin sodium salt (Regis Technologies) were anesthetized and euthanized. Organs (lungs, liver, spleen, heart, kidney, and brains) were collected, organized on a sheet, and imaged for chemiluminescence in an IVIS Spectrum machine (Revvity). Revvity LivingImage software (version 4.5.5) was used to analyze images.

### Lung/Liver Flow Cytometry

Naive mice were intravenously injected with either αPECAM-Fab DNA-LNPs formulated with 0.3% DSPE-PEG-Fluor 488 (Broad Pharm), to track particle delivery, or normal αPECAM-Fab DNA-LNPs loaded with pDNA encoding mCherry (Aldevron). Either 30 minutes (fluorescent LNPs) or 24 hours (mCherry LNPs) after treatment, mice were euthanized and transcardially perfused with PBS to clear blood and preserve tissue. The lungs and liver of the mice were collected and triturated before being incubated in a digestive solution of 2 mg/mL type 1 collagenase (Gibco) and 100μL of 2.5 mg/mL DNAse (Roche) for 45 min to prepare a single-cell suspension. After incubation, the cells were strained through a 70μm cell strainer (Sycamore Life Science) and washed with PBS. The supernatant was discarded, and ACK lysis buffer (Gibco) was added for 5 min on ice to lyse any remaining red blood cells (RBCs). After RBC lysis, an automated cell counter (Countess, Thermo Fisher) was used to achieve a final cell concentration of 1 × 10^6^ cells/mL for each sample. Suspensions were washed with Fluorescence-activated Cell Sorting (FACS) buffer, consisting of PBS, 1% Fetal Bovine Serum (FBS, Thermo Fisher Scientific), and 1 mM Ethylenediaminetetraacetic Acid (EDTA, Invitrogen) and incubated with anti-CD16/CD32 monoclonal antibodies (Invitrogen) for 15 minutes to prevent nonspecific binding of antibodies on Fc-receptors. Following washing, suspensions were incubated for another 30 mins with an antibody cocktail consisting of brilliant ultra violet (BUV) 395 anti-mouse CD45, clone: 30-F11 (BD Biosciences), phycoerythrin (PE)/cyanine7 anti mouse-CD64, clone: FcγRI (BioLegend), and alexa fluor 700 anti-mouse Ly6G, clone: Gr-1 (BioLegend) to identify immune cell subpopulations and allophycocyanin (APC) anti-mouse CD31 (BioLegend) and brilliant violet 711 (BV711) anti-mouse Ep-CAM, clone:CD326 (BioLegend) for endothelial and epithelial cells. After antibody incubation, the cells were fixed with 4% paraformaldehyde (PFA, Electron Microscopy Sciences) before analysis on a flow cytometer (LSR Fortessa, BD BioSciences). Gating parameters for analysis are illustrated in **Figure S11**. The flow cytometry results were analyzed using FlowJo™ Software v10.10 (BD Life Sciences).

### Brain Flow Cytometry

Naive mice were intravenously injected with αVCAM-mAb DNA-LNPs loaded with pDNA encoding mCherry (Aldevron). 24 hours after treatment, mice were euthanized and transcardially perfused with Hank’s Balanced Salt Solution (HBSS without Ca2+ and Mg2+, Thermo Fisher Scientific) to clear blood and preserve tissue. The brains of the mice were collected and placed in a small weighing boat with 2mL HBSS and passed through a 18g needle, then 21g needle 2-3 times until evenly homogenized. After homogenization, the cells were strained through a 100μm cell strainer (Sycamore Life Science) and washed with HBSS, using a black rubber stopper of 5mL syringe to press the homogenate through the filter. Cells were spun down at 280g for 5 minutes at 4C, the supernatant was discarded, and cells were resuspended in 1mL of dispase digestion buffer (100μL dispase solution, Theromo Fisher scientific, in 2mL HBSS) and incubated in a rotator at 37C for 1 hour. Once again, the cells were strained through a 70μm cell strainer (Sycamore Life Science) and washed with 1mg/mL DNAse (Roche) in HBSS before being spun down at 280g for 5 minutes at 4C. Cells were resuspended in 4mL of density gradient (1mL NaCl, 9mL Percoll (Thermo Fisher Scientific) and 30mL HBSS) and spun at 520g for 20 minutes at 4C. The supernatant was discarded, and ACK lysis buffer (Gibco) was added to cells for 30 seconds on ice to lyse any remaining RBCs. After RBC lysis, an automated cell counter (Countess, Thermo Fisher) was used to achieve a final cell concentration of 1 × 10^6^ cells/mL for each sample. Suspensions were washed with Fluorescence-activated Cell Sorting (FACS) buffer, consisting of PBS and 2% Fetal Bovine Serum (FBS, Thermo Fisher Scientific) and incubated with anti-CD16/CD32 monoclonal antibodies (Invitrogen) for 15 minutes to prevent nonspecific binding of antibodies on Fc-receptors.

Following washing, suspensions were incubated for another 30 mins with an antibody cocktail consisting of brilliant ultra violet (BUV) 395 anti-mouse CD45, clone: 30-F11 (BD Biosciences), phycoerythrin (PE)/cyanine7 anti mouse-CD64, clone: FcγRI (BioLegend), and alexa fluor 700 anti-mouse Ly6G, clone: Gr-1 (BioLegend) to identify immune cell subpopulations and allophycocyanin (APC) anti-mouse CD31 (BioLegend) and brilliant violet 711 (BV711) anti-mouse Ep-CAM, clone:CD326 (BioLegend) for endothelial and epithelial cells. After antibody incubation, the cells were fixed with 4% paraformaldehyde (PFA, Electron Microscopy Sciences) before analysis on a flow cytometer (LSR Fortessa, BD BioSciences). All single cells were included in our brain flow analysis, so no gating beyond forward versus side scatter was applied (first panel of **Figure S11**). The flow cytometry results were analyzed using FlowJo™ Software v10.10 (BD Life Sciences).

### Histology

Either naive BALB/c or B6.Cg-Gt(ROSA)26Sortm6(CAG-ZsGreen1)Hze (Ai6 for short) mice were intravenously injected with DNA-LNPs (conjugated to either αPECAM Fabs for lung histology, αVCAM mAbs for brain histology, or IgG as an untargeted control loaded with pDNA (Aldevron) encoding mCherry 4 hours or Cre recombinase 24 hours prior to perfusion. For brain histology, mice were perfused through the left ventricle (after cutting the right atrium) with 15mL 1x PBS followed by 15mL 4% PFA (EMS). For lung histology, mice were perfused with 10mL 1x PBS through the right ventricle at 20cm H2O, the trachea was sutured shut, and the lungs were inflated with 4% PFA by injection into the trachea. Organs were then drop-fixed in 4% PFA for 16-24 hours at 4 degrees C and then cryo-protected in 30% sucrose (Neta Scientific) for at least 48 hours, embedded in OCT (Fisher), cryo-sectioned at 16um or 35um (in the case of lung confocal microscopy), mounted on Superfrost Plus slides (Fisher), and frozen at −80 degrees C. Slides were then thawed at 60 degrees C, rehydrated in 1x PBS, and blocked for one hour at RT in blocking buffer - 90% 1x PBS, 9.5% donkey serum (Sigma), 0.5% Triton X-100 (Sigma). After blocking, slides were incubated with primary antibodies (described below) in staining buffer (98.5% 1x PBS, 1% donkey serum, 0.5% Triton X-100) overnight at 4 degrees C. In the morning, slides were washed with 1x PBS and incubated with secondary antibodies (described below) for one hour at RT in staining buffer. Slides were washed once more and coverslipped with DAPI mounting media (EMS) before imaging. Controls at identical exposure settings for both lung and brain histology are shown in **Figure S7**.

Antibodies used: Primaries included Goat Anti-CD31(Biotechne, 1:200); Rabbit Anti-Iba1 (Fujifilm, 1:600); Rat Anti-GFAP (Agilent, 1:400); Guinea pig Anti-S100B (BD Biosciences, 1:400); Chicken polyclonal Anti-mCherry (Abcam, 1:200); and Mouse Anti-NeuN(Agilent, 1:200). Secondaries included Donkey anti-goat IgG 488, 647 (ThermoFisher, 1:500); Donkey anti-rabbit IgG 594, 647 (ThermoFisher, 1:500); Donkey anti-rat IgG 488 (ThermoFisher, 1:500) Donkey anti-chicken 647 (ThermoFisher, 1:500); Donkey anti-mouse IgG 488 (ThermoFisher, 1:500); Donkey anti-guinea pig 647 (BD biosciences, 1:500). Microscopes used: BZ-x800 widefield fluorescent microscope and Andor benchtop confocal microscope. Fiji and Imaris softwares used for image processing and analysis.

Quantification: To quantify the percent of expressing cells with targeted vs untargeted DNA-LNPs, we imaged 3 regions of interest (ROIs) at 20X magnification for cerebellar and pericerebellar regions for *n* = 2 mouse sagittal brain sections per condition for Cre DNA-LNPs (αVCAM vs IgG) and *n* = 3 mouse sagittal brain sections per condition for mCherry DNA-LNPs (αVCAM vs IgG). We performed quantification using Fiji software and a custom macro that executed the following logic: for both the DAPI and reporter channel of interest (488 for Cre and 594 for mCherry) convert to 8bit, threshold (kept the same for all Cre or all mCherry images), make binary, count and save DAPI masks, calculate percent overlap of DAPI masks on merge with reporter channel masks. Values reported as % reporter positive DAPI positive cells. Dot render generation implemented Fiji software to threshold, create masks, and circularize masks into dot signals which were overlaid onto the composite image for improved visualization as done previously^44^.

### Reverse transcription quantitative polymerase chain reaction (RT-qPCR)

Naive BALB/c mice received intravenous injections of either αVCAM DNA-LNPs or IgG DNA-LNPs, each loaded with plasmid DNA encoding mCherry (Aldevron). 24h post-dose, mice were euthanized and transcardially perfused with 1x PBS. Identical regions of brain (one sample from cortex, one sample from cerebellum) weighing < 30mg were collected, homogenized, and lysed in 1mL Buffer RLT containing 20µL of 2M dithiothreitol (DTT). Total RNA was isolated using the RNeasy Mini Kit (Qiagen) according to the manufacturer’s Quick-Start protocol for tissues. To remove genomic DNA, we performed on-column DNase digestion using the RNase-Free DNase set. RNA concentration was quantified by NanoDrop and stored on ice for immediate cDNA synthesis. cDNA was generated from 200-400ng total RNA using the Thermo Scientific Verso cDNA Synthesis Kit with a 3:1 blend of random hexamers:anchored oligo-dT, following the kit instructions. qPCR was performed with SYBR Green chemistry on a 384-well real-time instrument (insert brand of instrument). Each 10µL reaction contained 6µL SYBR Green reagent, 20µM forward primer, 20µM reverse primer, and 4ng cDNA (or a fixed input volume applied uniformly across samples). Cycling used a standard program in QuantStudio: 95C for 20 sec, then 40 cycles of 95C 3s and 60C 30s with plate read. Primers targeted mCherry (gene of interest) and β-actin and GAPDH (housekeepers).

Quantification: Data were analyzed with automatic, uniform baseline/threshold settings. ΔCt was defined as Ct(mCherry) − Ct(geometric mean of β-actin and GAPDH). For relative quantification, IgG-LNP served as the calibrator: ΔΔCt = ΔCt(sample) − mean ΔCt(IgG); fold-change = 2^-ΔΔCt^, known as the comparative Ct method. Consistent with best practice, statistics were performed on ΔCt (or equivalently log_2_[fold-change]), while 2^−ΔΔCt^ (fold change) was plotted for interpretability.

### Serum Multiplex

For plasma collection, mice treated with 5μg αPECAM-Fab DNA-LNPs LNPs were euthanized by terminal blood collection using EDTA syringes through the inferior vena cava. Opening of the major body cavity and subsequent thoracotomy was performed as a secondary measure of euthanization. Blood was centrifuged at 1,000g for 10 minutes at room temperature, and then plasma supernatant was collected and stored at −80C. Cytokine measurements were carried out on plasma (2x diluted) with a LEGENDplex 13-plex Mouse Inflammation Panel (BioLegend) according to the manufacturer’s instructions.

### Weight Collection and Pulse Oximetry

Naive BALB/c mice were split up into either a naive (control) or treatment group, which received 5μg injection of αPECAM-Fab DNA-LNPs. Prior to measurement with pulse oximetry (MouseOx Plus), mice were placed in a chamber with 2% isoflurane for induction anesthesia. Following induction, mice were moved to an anesthesia nose cone, where (1) a pulse oximetry collar was placed and (2) anesthesia was titrated to maintain between 150-200 breaths per minute (maintenance isoflurane 0.5-1%). Respiratory rate and oxygen saturation was sampled at high frequency. Epochs with acceptable signal quality were used for analysis by obtaining values from artifact-free windows of >10s (which were averaged for analysis). Daily weights were obtained starting from time of injection (*t* = 0h) and every 24h thereafter.

### Statistical Analysis

Statistical analyses were performed using GraphPad Prism 10 (GraphPad Software). All data include n = 3 (biological replicates) and represent mean ± SEM, unless otherwise specified; * = p < 0.05, ** = p < 0.01, *** = p < 0.001, **** = p < 0.0001, ns = not significant. The statistical tests used for each experiment are described in each corresponding figure caption.

## Data Availability

All data that support the findings of this study are available from the corresponding author upon request.

## Acknowledgments

N.M. and T.V.B. contributed equally to this work. Research reported in this publication was supported by the National Science Foundation under Cooperative Agreement DBI – 2400327 (to T.V.B. and V.R.M.), the National Center for Advancing Translational Sciences of the National Institutes of Health-KL2TR001879 (to E.E.), National Library of Medicine under the National Institutes of Health-R01NS131279 (For O.A.M.C.), the National Institute of Health-1DP5OD036159 (for M.L.B.), R01-HL-153510 (J.S.B.), -HL160694 (J.S.B.), -HL164594, (J.S.B.), -HL-157189 (J.S.B.), -HL-155106 (V.R.M.) and the American Heart Association under grant 24PRE1195406 (to M.N.P.). The flow cytometry data for this manuscript were generated in the Penn Cytomics and Cell Sorting Shared Resource Laboratory at the University of Pennsylvania (RRID:SCR_022376). Penn Cytomics is partially supported by the Abramson Cancer Center NCI Grant (P30 016520). The content is solely the responsibility of the authors and does not necessarily represent the official views of the National Institutes of Health.

