## Supplemental Data for "Targeting DNA-LNPs to Endothelial Cells Improves Expression Magnitude, Duration, and Specificity"

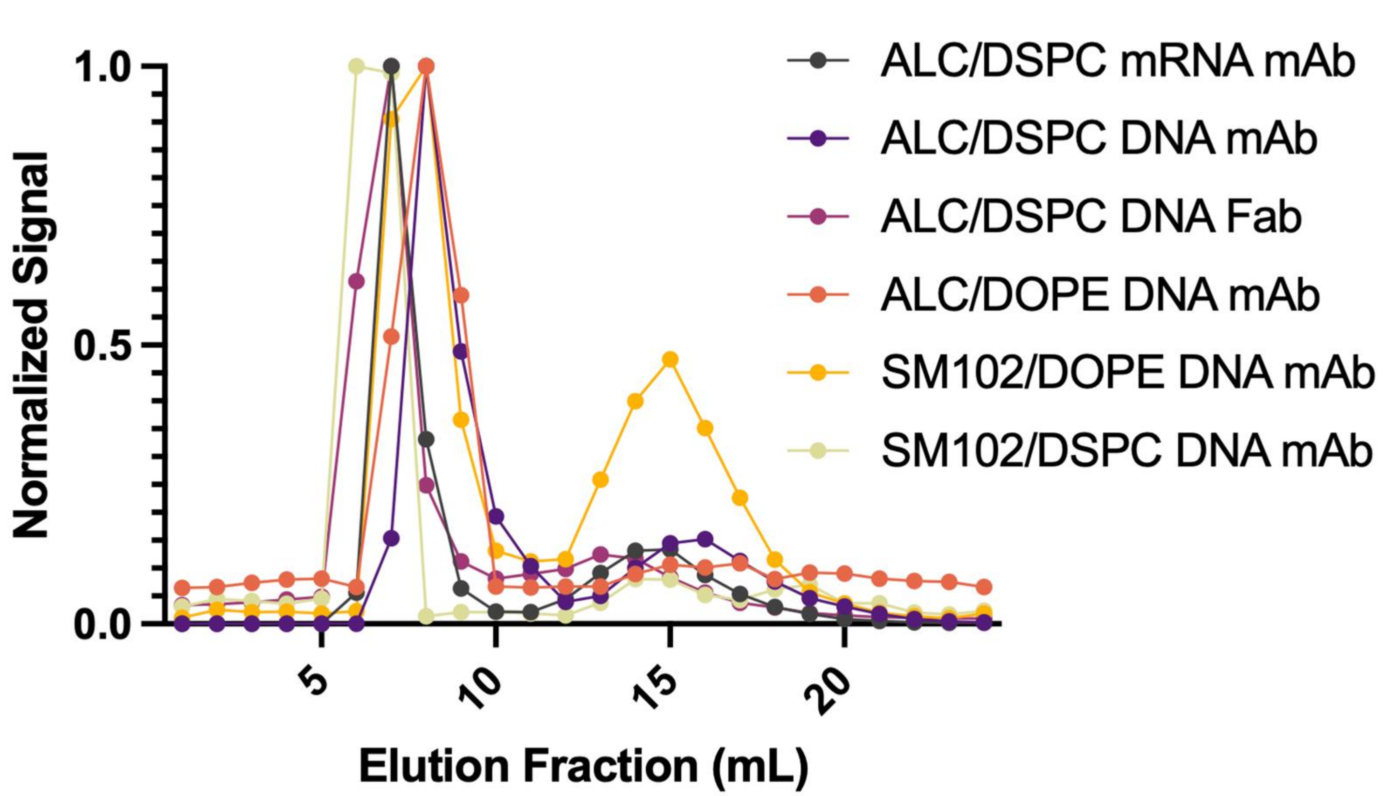


| Formulation | Conjugation Efficiency |
|---|---|
| ALC/DSPC mRNA mAb | 85.7% |
| ALC/DSPC DNA mAb | 84.3% |
| ALC/DSPC DNA Fab | 83.8% |
| ALC/DOPE DNA mAb | 80.9% |
| SM102/DSPC DNA mAb | 80.3% |
| SM102/DOPE DNA mAb | 68.5% |

**Supplemental Figure 1. Elution profiles of different formulations of antibody conjugated DNA-LNPs.** Elution curves of different DNA-LNP formulations conjugated to various $^{125}$I-labeled targeting moieties (mAb = αPECAM monoclonal antibody; Fab = αPECAM fragment antigen binding region). Using Sepharose size exclusion chromatography (SEC), we anticipate antibody conjugated LNPs to elute in the first peak (between fractions 6-9), with unbound antibodies in subsequent peaks. Elution peaks were normalized by taking the raw CPM signal from each fraction (measured via Gamma counting) and dividing it by the largest CPM value from all the fractions. We used these curves to generate **Table S1**, a summary of conjugation efficiencies for each DNA-LNP formulation.

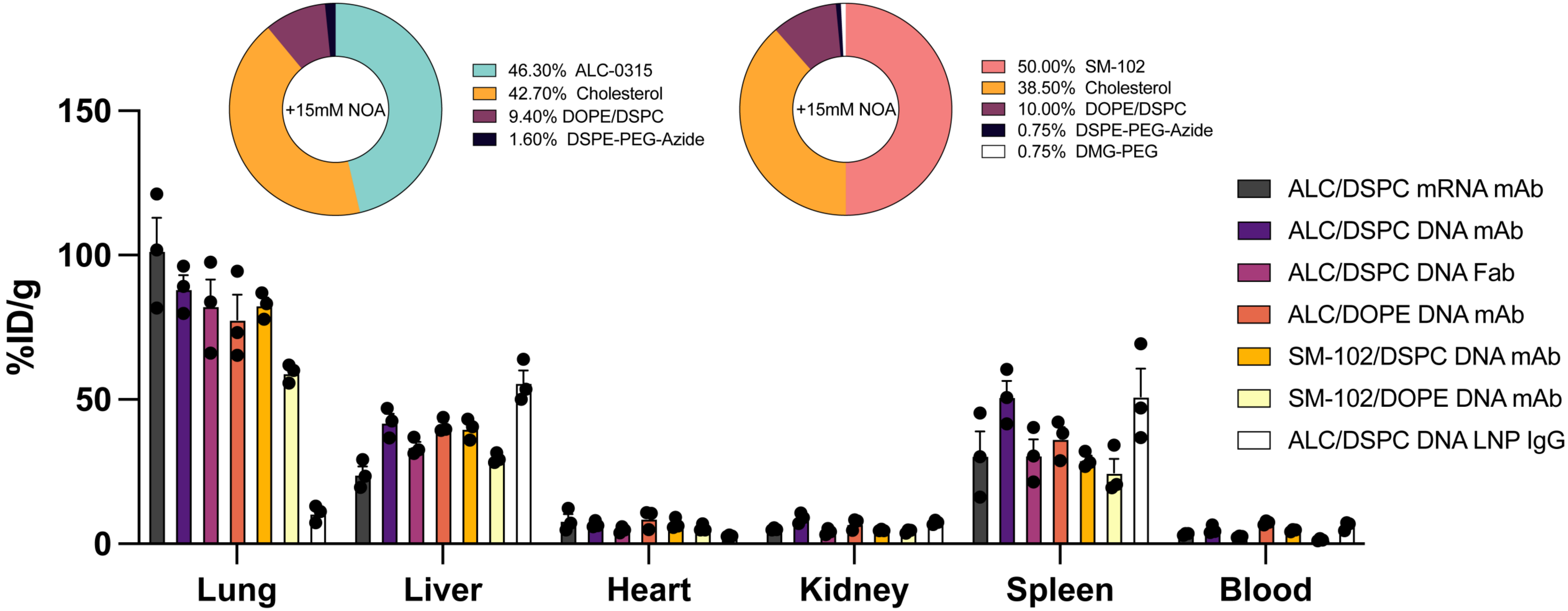


**Supplemental Figure 2. Representative biodistribution of additional formulations of DNA-LNPs.** Biodistribution study via radiotracing of different formulations of $^{125}$I-labeled DNA-LNPs conjugated to various targeting moieties (mAb = αPECAM monoclonal antibody; Fab = αPECAM fragment antigen binding region; IgG = immunoglobulin G control). Detailed lipid formulations are shown in pie charts above. %ID/g represents the percent of total injected dose detected in each organ normalized by respective organ mass.

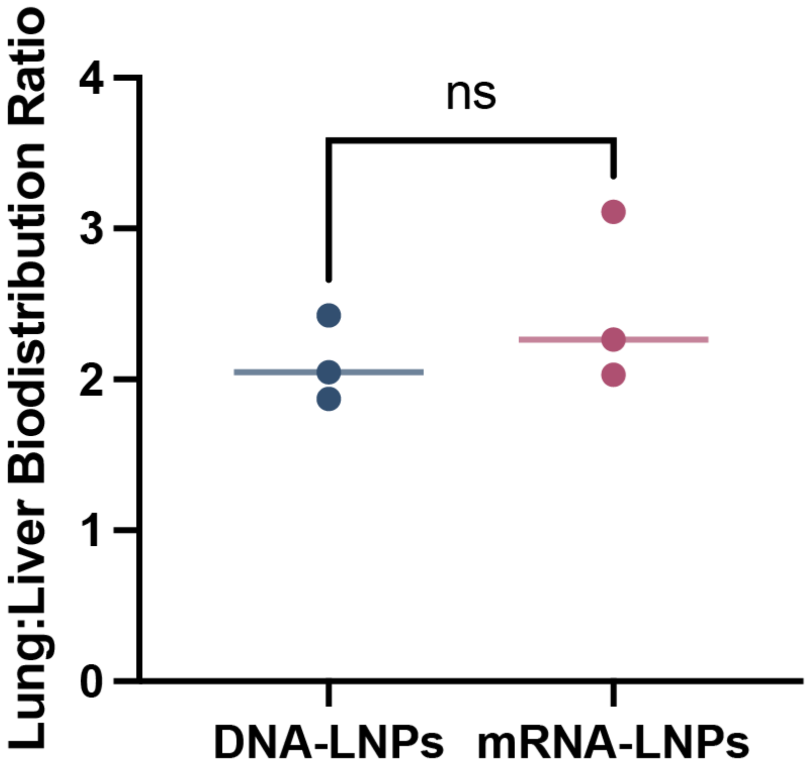


**Supplemental Figure 3. Comparison of Biodistribution Ratios Between mRNA- and DNA-LNPs.** Biodistribution study via radiotracing mRNA- or DNA-LNPs conjugated to αPECAM mAbs (50 mAbs/LNP) and untargeted $^{125}$I-IgG to track particle distribution (5 mAbs/LNP) shows no significant difference in lung-to-liver biodistribution ratios, which is represented as the %ID/g measured in the lungs divided by the %ID/g measure in the liver. %ID/g represents the percent of total injected dose detected in each organ normalized by respective organ mass.

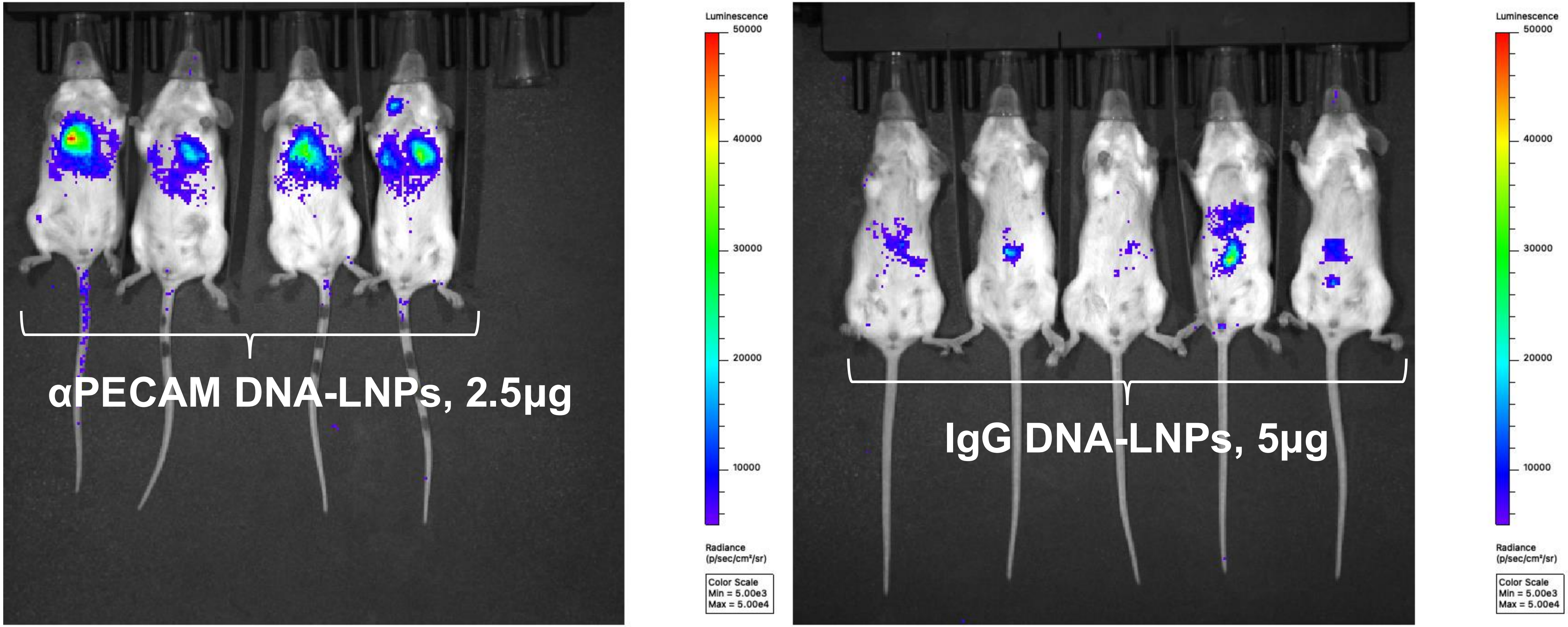


**Supplemental Figure 4. IVIS images of 2.5μg αPECAM DNA-LNPs and 5μg untargeted IgG DNA-LNPs.** Representative IVIS images of αPECAM DNA-LNP-treated BALB/c mice dosed with 2.5μg luciferase pDNA (left) and untargeted IgG DNA-LNP-treated mice dosed with 5μg luciferase pDNA (right) at 1d post-injection.

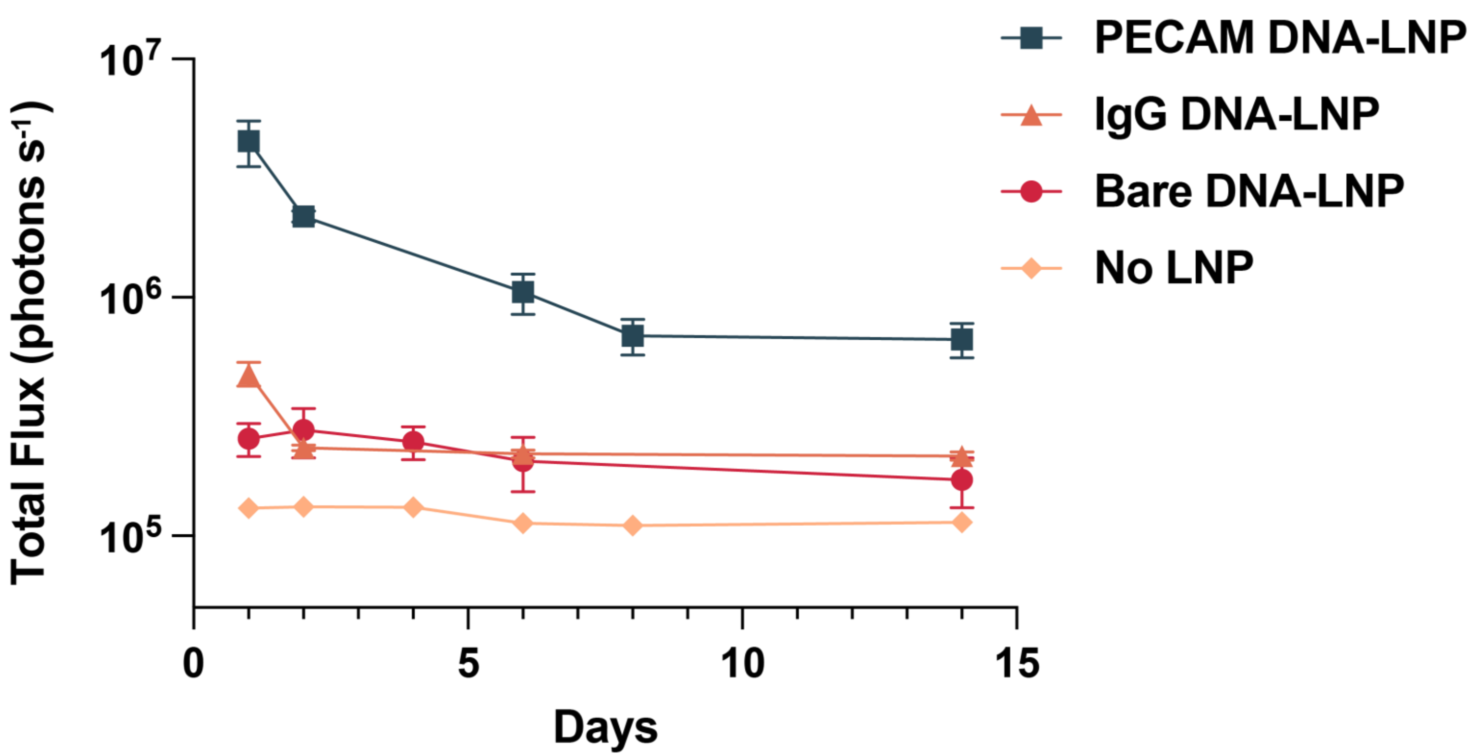


**Supplemental Figure 5. Quantified *in vivo* luminescence for untargeted IgG and bare DNA-LNPs.** Quantified in vivo luminescence for anti-PECAM (blue), bare (red), and untargeted IgG (orange) DNA-LNPs compared with untreated, luciferin-injected control mice (yellow). Total luminescence (represented as total flux in photons $s^{-1}$) in the area of visible signal from IVIS images over time for mice treated with 5μg of luciferase pDNA loaded into bare or untargeted IgG-conjugated LNPs. The expression is substantially lower than the total expression in mice treated with 5μg of αPECAM DNA-LNPs, but is higher than baseline, untreated expression.

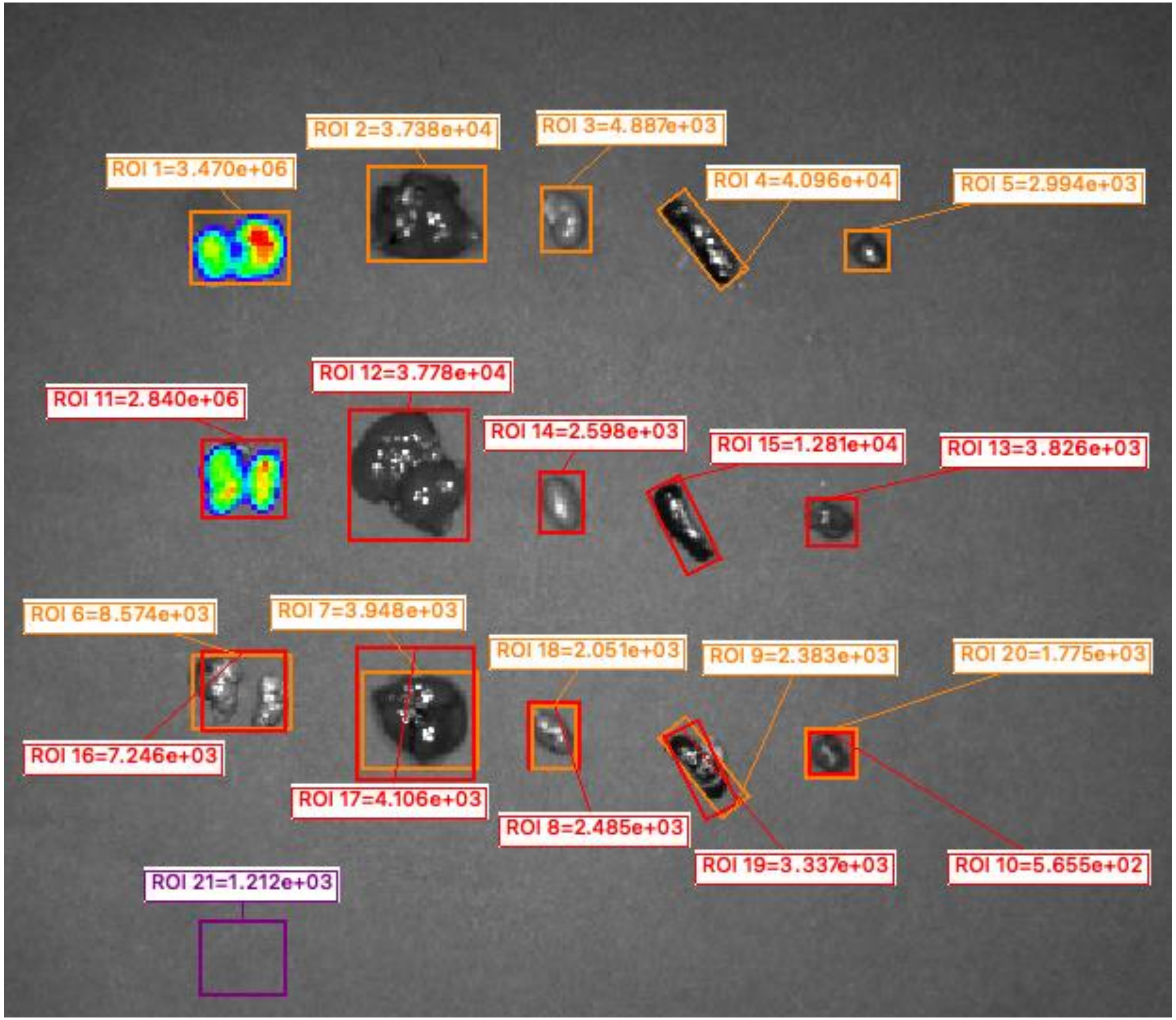


**Supplemental Figure 6. Quantification Method for *Ex Vivo* Bioluminescence.** Image depicting quantification method used for calculating *ex vivo* luminescence (in photons $s^{-1}$) at various regions of interest (ROIs). Briefly, ROIs are first drawn around organs to capture entire organ while minimizing background captured. Next, size-matched ROIs are drawn around control organs (from luciferin-injected mice) and quantified. Finally, control values (considered background) are subtracted from treatment values. A background ROI quantification is shown (purple) to underscore how low plate background is ($1.212 \times 10^3$) relative to sample measurements (*i.e.*, $3.47 \times 10^6$).

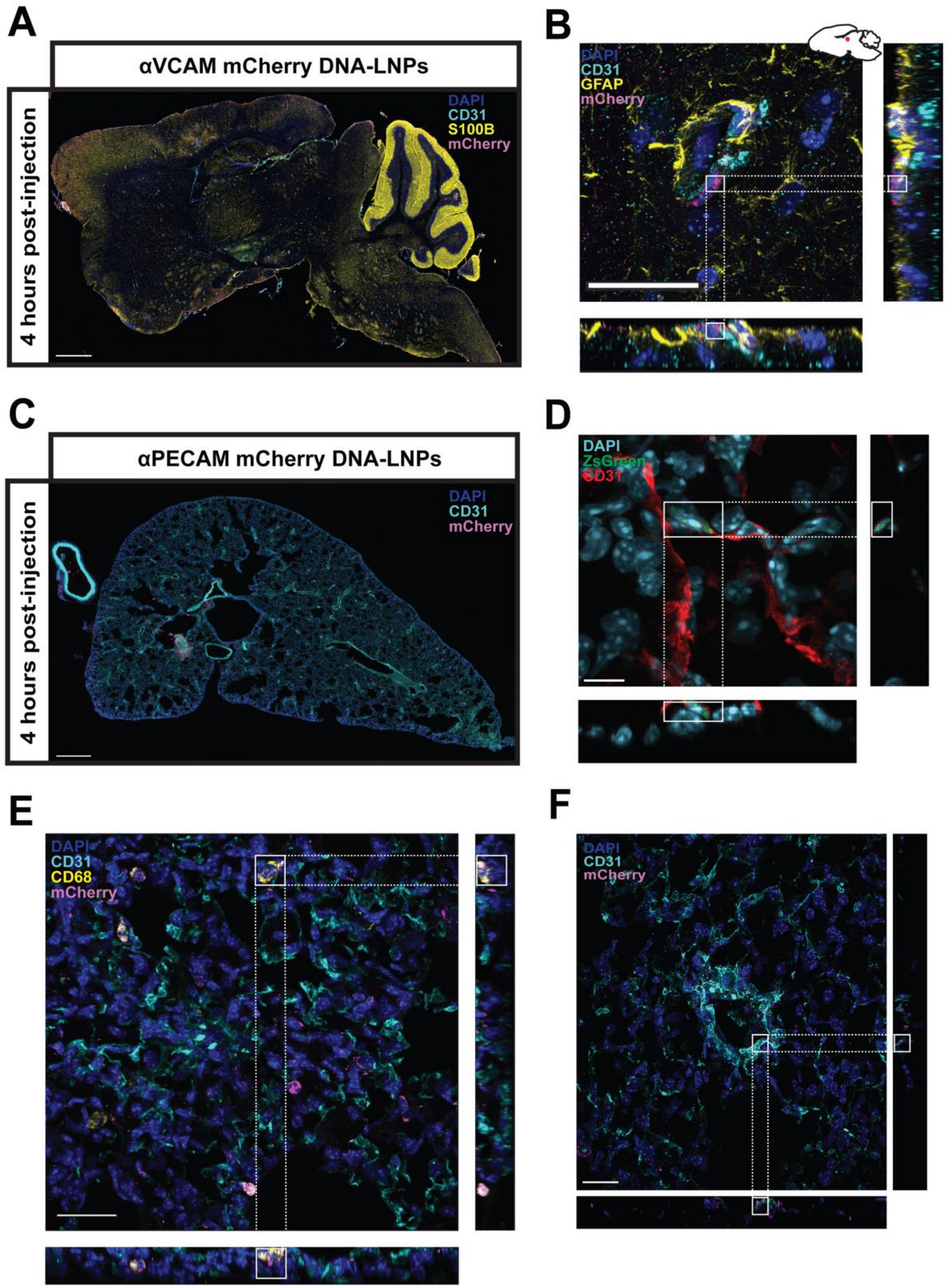


**Supplemental Figure 7. Immunofluorescence of Lungs and Brain demonstrating targeted transgene expression after treatment with conjugated pDNA-LNPs.** (A) Immunofluorescence sagittal brain section of a naive BALB/c mouse 4 hours after treatment with αVCAM DNA-LNPs loaded with mCherry pDNA. Merge depicts DAPI (blue), PECAM-1 (cyan), mCherry (magenta), and the astrocyte marker S100B (yellow). Scale bar, 1mm. (B) 60X image with orthogonal views highlighting endothelial cells (CD31, cyan), astrocytes (GFAP, yellow), and mCherry signal (magenta) (scale bar, 30µm). (C) IF transverse lung section (20X resolution) of a naive BALB/c mice 4 hours after treatment with αPECAM Fab DNA-LNPs loaded with mCherry pDNA. Merge shows DAPI (blue), PECAM-1 (cyan), and mCherry (magenta) (scale bar, 1mm). (D) 60X image with orthogonal views demonstrate ZsGreen expression in endothelial cells (CD31, red) lining the blood vessels of the pulmonary vasculature 24 hours after injection with αPECAM Fab DNA-LNPs loaded with Cre pDNA (scale bar, 50µm). (E) 60X image with orthogonal views of BALB/c lung 4 hours after treatment with αPECAM Fab DNA-LNPs loaded with mCherry pDNA. Merge shows DAPI (blue), CD31 (cyan), CD68 (yellow), and mCherry (magenta) highlighting positive expression in myeloid cells (scale bar, 50µm). (F) 60X image with orthogonal views demonstrates mCherry expression in endothelial cells (CD31, cyan) lining the blood vessels of the pulmonary vasculature 4 hours after injection with αPECAM Fab DNA-LNPs loaded with Cre pDNA (scale bar, 50µm).

**Supplemental Figure 8. Immunofluorescence of control IgG DNA-LNP brain expression 4h and 24h post-treatment.** (A & B) Cerebellar and pericerebellar immunofluorescence brain sections of an Ai6 mouse 24 hours after treatment with IgG DNA-LNPs loaded with Cre pDNA. Merge depicts DAPI (blue), S100B (white), and ZsGreen (green), and the astrocyte marker S100B (yellow). Scale bar, 100µm. (C & D) Cerebellar and pericerebellar immunofluorescence brain sections of a naive BALB/c mouse 4 hours after treatment with IgG DNA-LNPs loaded with mCherry pDNA. Merge depicts DAPI (blue), NeuN (yellow), CD31 (cyan), and mCherry (pink). Scale bar, 100µm.

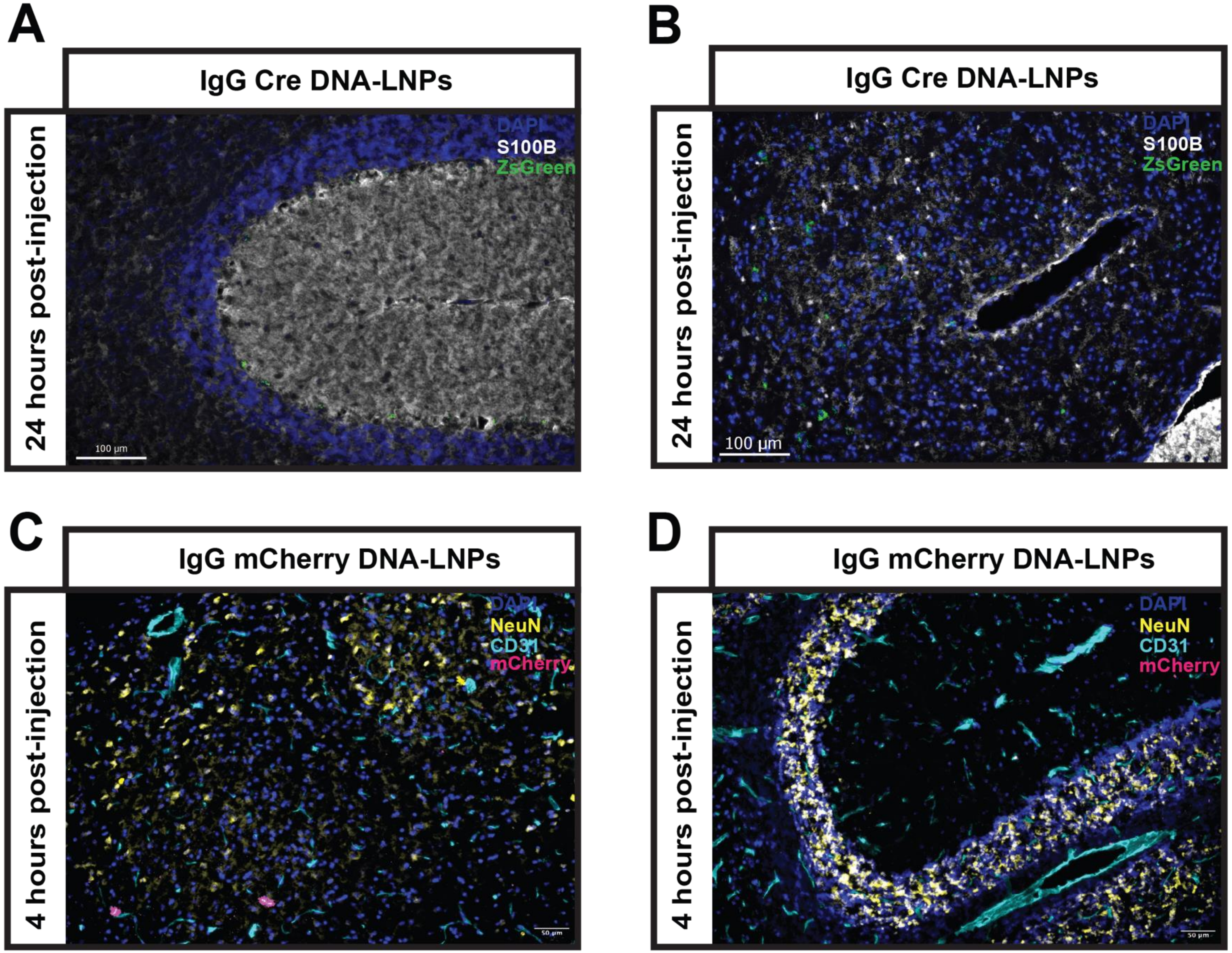

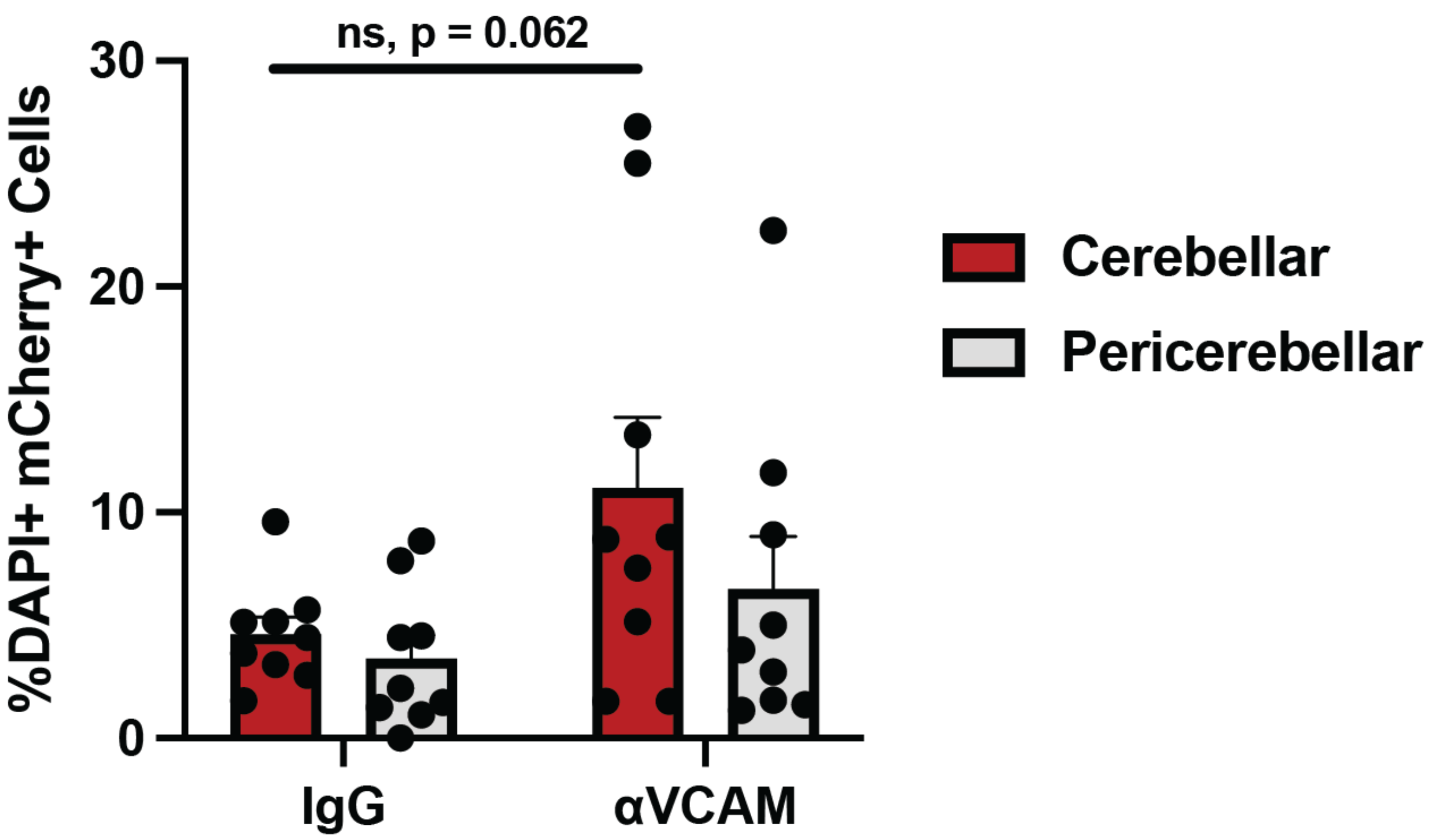


**Supplemental Figure 9. Quantification of IgG and VCAM mCherry DNA-LNP immunofluorescence.** At 4h post-treatment with either IgG or VCAM mCherry DNA-LNPs, brains were harvested for immunofluorescence. Quantification of cerebellar and pericerebellar brain slices with an unbiased macro shows no statistically significant differences in 4h expression, though the data appear to trend toward significance.

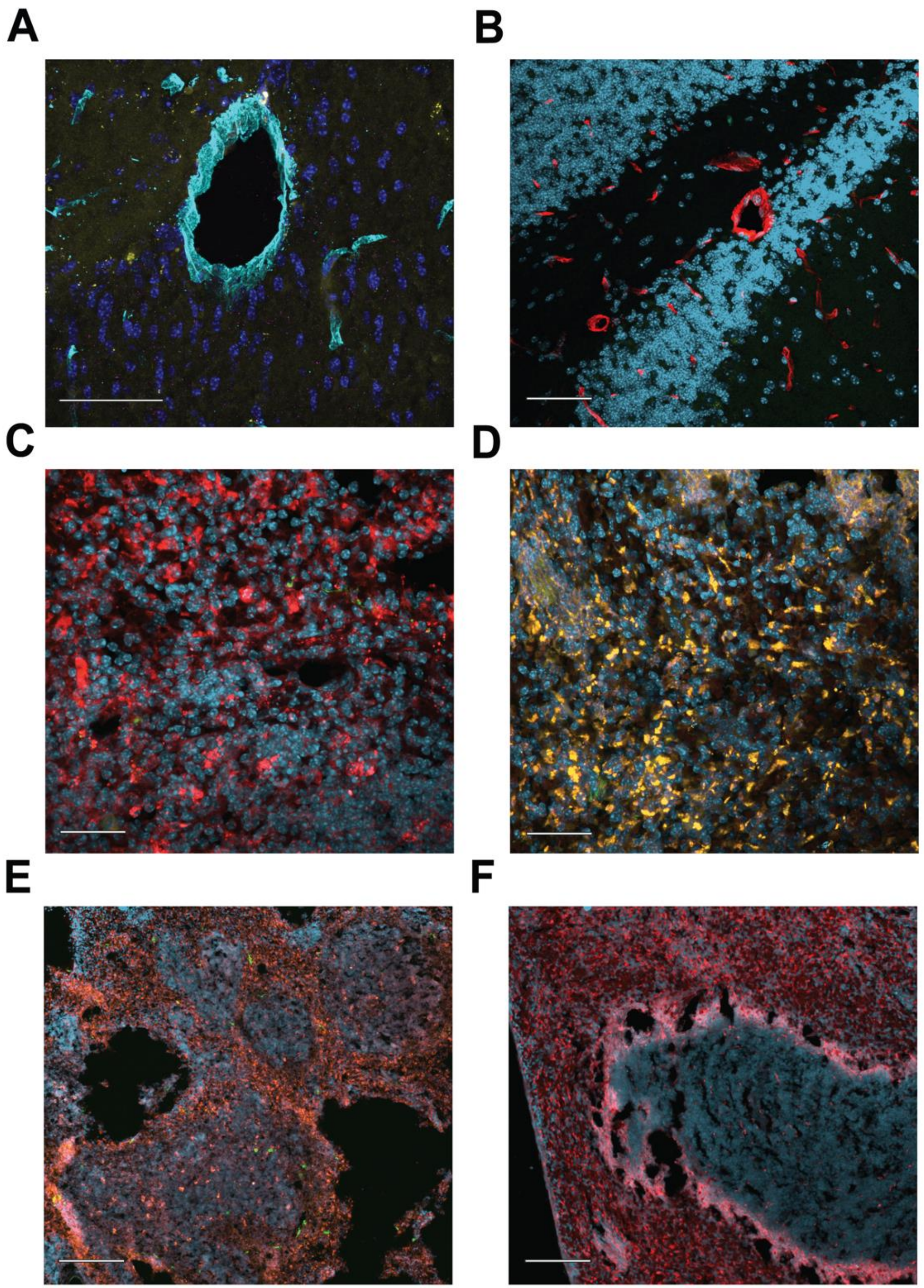


**Supplemental Figure 10. Additional Immunohistochemistry controls.** (A) 60X representative image of uninjected BALB/c mouse brain taken at the same exposure settings as brain tissue from mice treated with αVCAM DNA-LNPs loaded with mCherry pDNA with merge showing CD31 (cyan), no primary antibody with anti-rat 488 secondary antibody (yellow), anti-mCherry staining (magenta), and DAPI (blue) (scale bar, 50μm). (B) 60X representative image of uninjected BALB/c mouse brain taken at the same exposure settings as brain tissue from Ai6 mice treated with αVCAM DNA-LNPs loaded with Cre pDNA with merge showing CD31 (red), native 488 autofluorescence (green), and DAPI (cyan) (scale bar, 50μm). (C and D) Representative 60X images of Ai6 (C) or BALB/c (D) splenic tissue illustrating ZsGreen or native 488 autofluorescence (green), CD68 (red), and DAPI (cyan) (scale bar, 50μm). (E and F) Representative 20X images of Ai6 (E) or BALB/c (F) splenic tissue illustrating ZsGreen or native 488 autofluorescence (green), no primary antibody anti-Rabbit 594 nonspecific staining (red), and DAPI (cyan) (scale bar, 100μm).

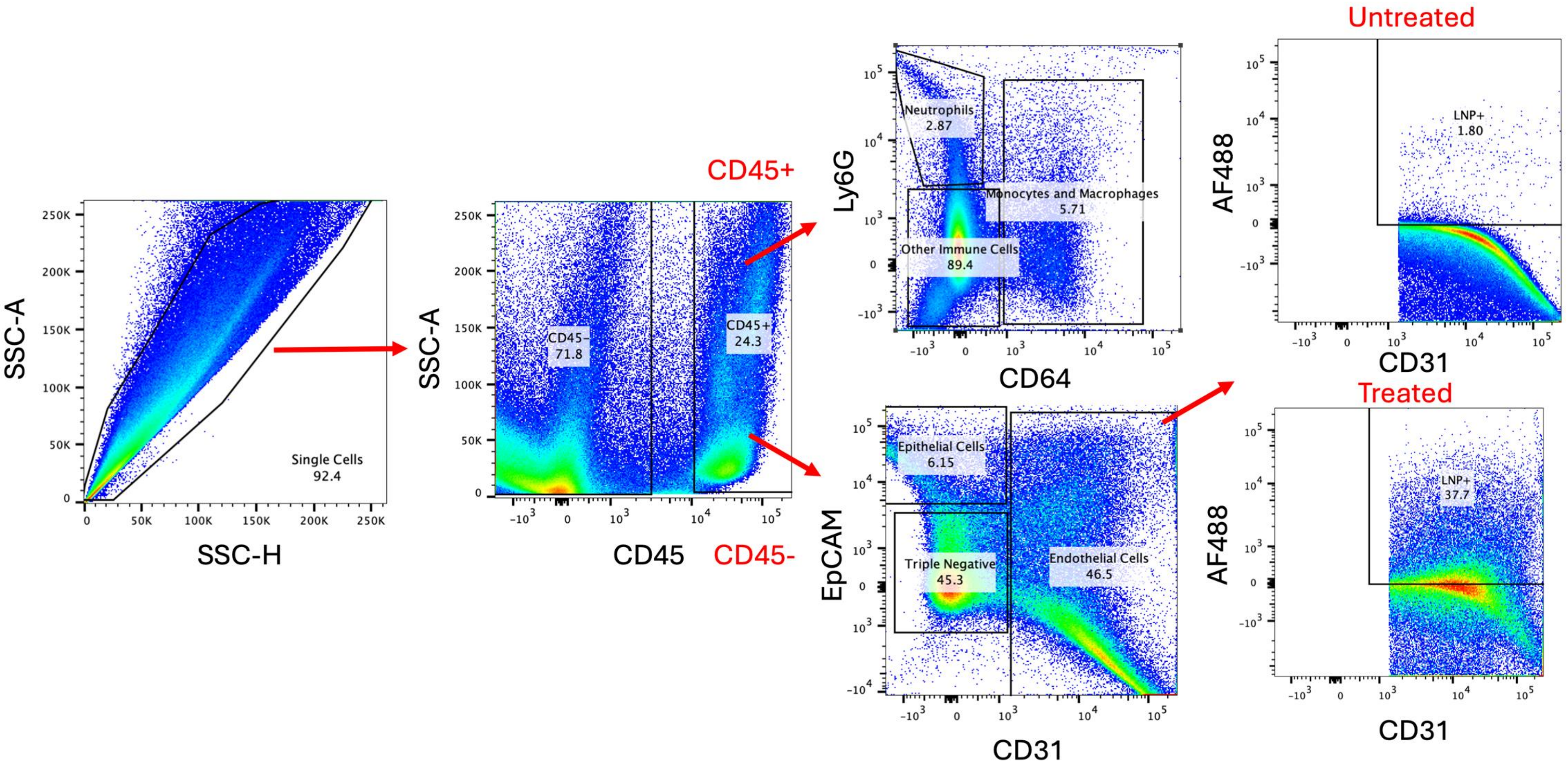


**Supplemental Figure 11. Gating strategy for flow cytometry.** To identify the cellular uptake of fluorescent DNA-LNPs in the lungs and liver, we first gated for single cells, then split the single cell population into CD45+ immune cells and CD45- non-immune cells. In the immune cell population, we identify neutrophils (Ly6G+), monocytes and macrophages (CD64+), and “other immune cells” (CD45+/Ly6G-/CD64-). In the non immune cell population, we identify endothelial cells (CD31+), epithelial cells (EpCAM+), and other “triple negative” cells (CD45-/CD31-/EpCAM-). To identify the LNP+ population, a fluorescence minus one (FMO) control was performed by surface staining for all cell types previously described in the organs of an untreated mouse. We set the upper threshold for a LNP+ cell based on the Alexa Fluor 488 signal of the FMO negative control (right side of figure). This same strategy was used when analyzing mCherry pDNA expression, as we set the upper threshold for an mCherry+ cell based on the mCherry signal of the untreated FMO control.

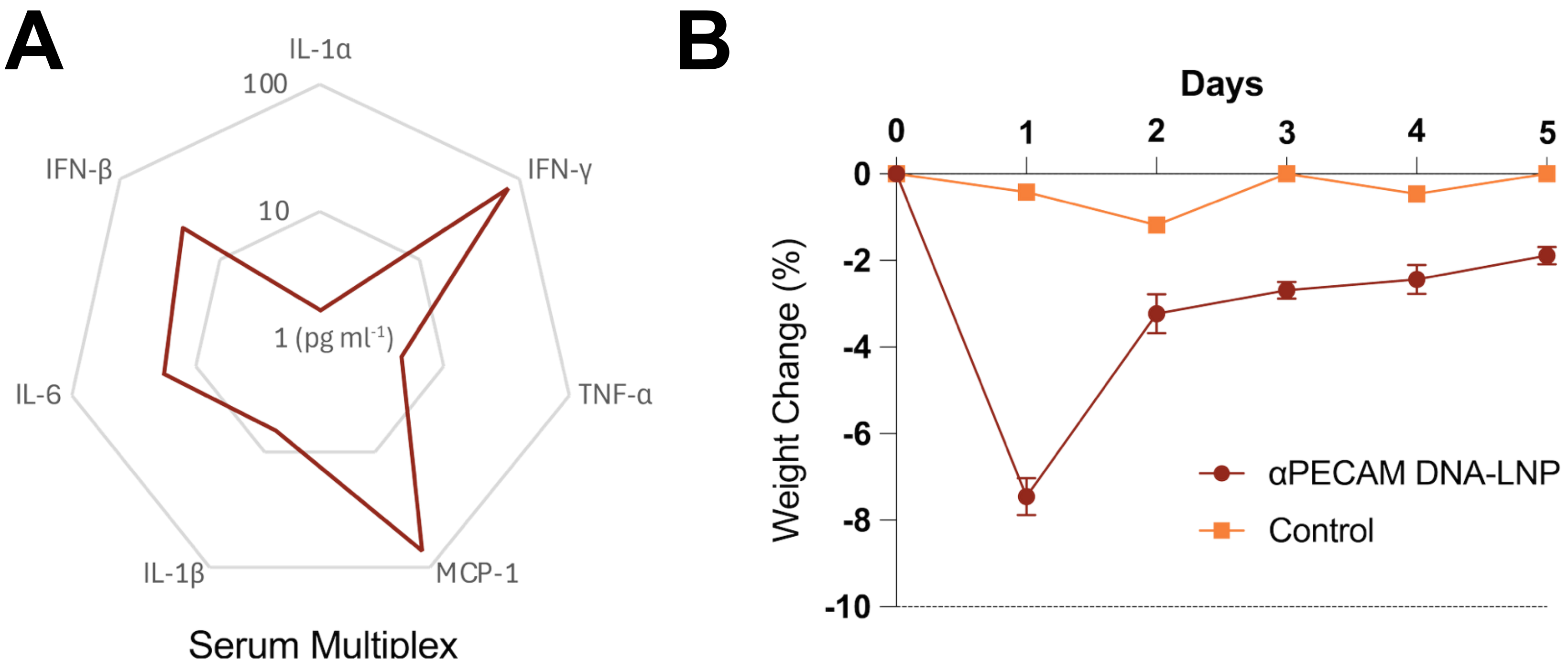


C

| Oxygen Saturation (%) | |
|---|---|
| t = 4h | 99.15 ± 0.06 |
| t = 24h | 99.03 ± 0.06 |

**Supplemental Figure 12. Toxicity Studies of Fab PECAM DNA-LNPs.** (A) Quantification of pro-inflammatory plasma cytokines 4-hours post 5 μg dose of αPECAM Fab DNA-LNPs. Notably, these levels are substantially lower than those reported in our initial study with untargeted DNA-LNPs. (B) Weight loss of naive (control) mice and mice treated with a 5 μg dose of αPECAM Fab DNA-LNPs. Treated mice have less than 10% weight loss (with no mortality). (C) Pulse oximetry measurements of mice treated with a 5 μg dose of αPECAM Fab DNA-LNPs at t = 4h and 24h post-injection. All measured oxygen saturations are above 99%.
